# PCSK9 deficiency promotes the development of peripheral neuropathy

**DOI:** 10.1101/2024.03.03.583154

**Authors:** Ali K. Jaafar, Aurélie Paulo-Ramos, Guillaume Rastoldo, Bryan Veeren, Cynthia Planesse, Matthieu Bringart, Philippe Rondeau, Olivier Meilhac, Gilles Lambert, Steeve Bourane

**Author notes:** **Corresponding Author: Steeve Bourane**. Université de la Réunion - INSERM U1188 - UMR DéTROI, UFR SANTE – TERRE SAINTE, 77 Avenue du Docteur Jean-Marie Dambreville, 97410 Saint-Pierre - La Réunion – France.

## Abstract

PCSK9 best-known and studied function is to induce the hepatic degradation of the low-density lipoprotein receptor (LDLR), thereby increasing the concentration of LDL-cholesterol (LDL-C) in the blood. Beyond its effects on LDL, recent studies have reported pleiotropic effects of PCSK9 notably in septic shock, vascular inflammation, viral infection, and cancer. While the functional and structural integrity of peripheral nerves are critically influenced by circulating lipids, the impact of PCSK9 in the peripheral nervous system is unknown. In this study, we investigated the consequences of PCSK9 deficiency on peripheral nerves. We found that PCSK9 deletion in mice leads to peripheral neuropathy characterized by a reduction of thermal and mechanical pain sensations. PCSK9 deficient mice also presented skin structural changes with a reduction of number of terminal nociceptive Schwann cells, Remak fiber axonal swelling, as well as hypomyelination of small nerve fibers. Interestingly, peripheral nerves of PCSK9 deficient mice presented an upregulation of the fatty acid transporter CD36 expression which correlated with an increase in nerve lipid contents and structural mitochondrial abnormalities. Our findings demonstrate that PCSK9 plays a critical role in the peripheral nerves by regulating lipid homeostasis, and its deficiency could lead to the development of symptoms related to peripheral neuropathy.

**Graphical abstract:** 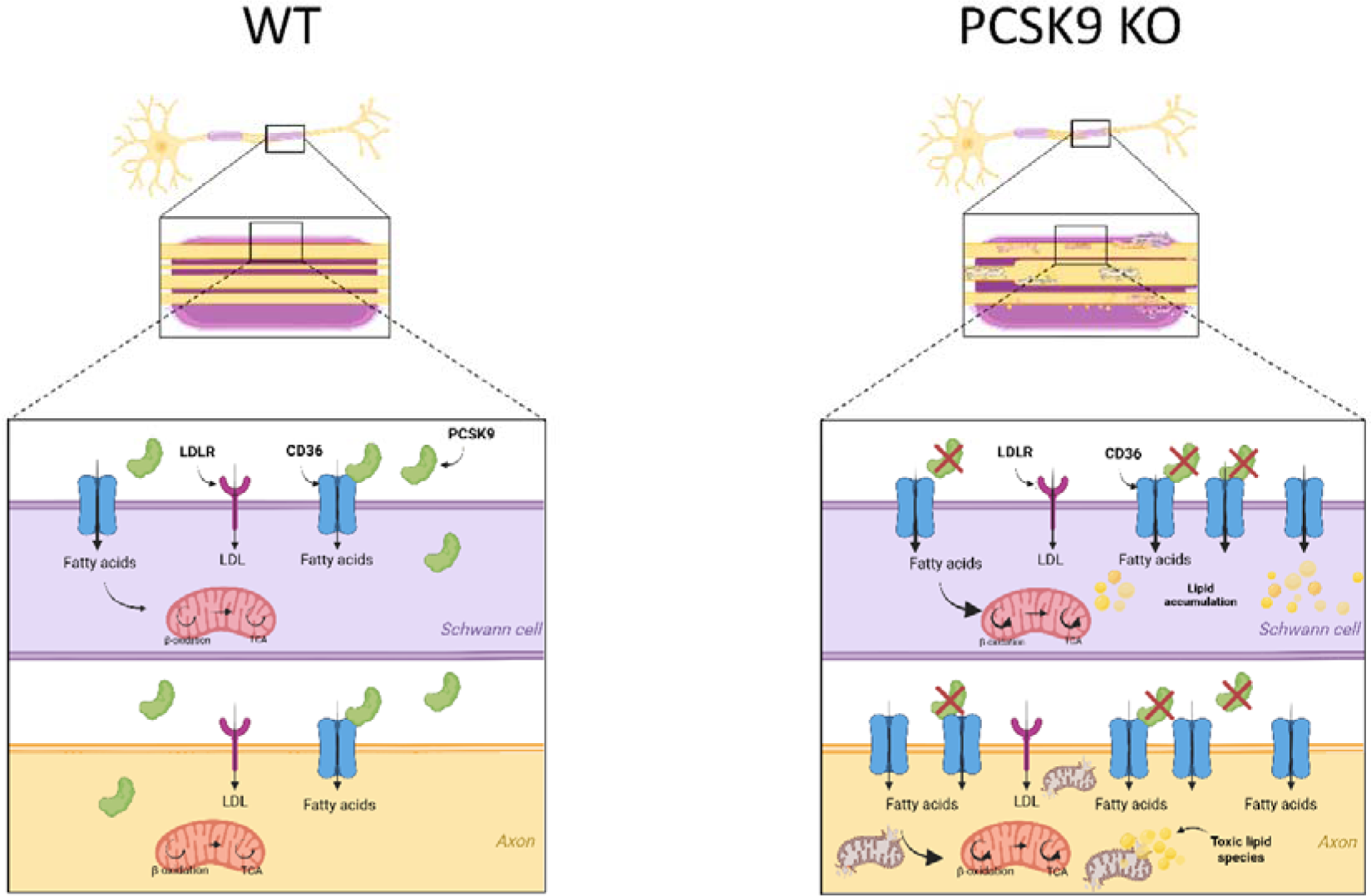

PCSK9 modulates nerve energy metabolism and health. Created with BioRender.com.

## Introduction

Proprotein convertase subtilisin/kexin type 9 (PCSK9) is a natural circulating inhibitor of the low-density lipoprotein receptor (LDLR) ^1^. It binds to the LDLR epidermal growth factor-A (EGFA) domain, and targets it for lysosomal degradation, which prevents LDLR recycling at the cell surface. As a result, PCSK9 increases the circulating levels of LDL-cholesterol (LDL-C). PCSK9 loss-of-function mutation carriers have reduced LDL-C levels and are protected against atherosclerotic cardiovascular diseases (ASCVD) ^2^, whereas PCSK9 gain-of-function mutation carriers present with familial hypercholesterolemia and are at high ASCVD risk ^3^. These observations led to the successful development of PCSK9 inhibitors, a new class of LDL lowering drugs, prescribed on top of statins, that have demonstrated their efficacy in large cardiovascular outcome trials ^4,5^. In addition, PCSK9 also reduces the abundance of LDLR related receptors, including the very-low density lipoprotein receptor (VLDLR), the LDLR related proteins 1 (LRP1) and 8 (LRP8, also known as apoER2), as well as the fatty acid transporter and scavenger receptor CD36 in different tissues ^1^.

PCSK9 is primarily expressed by the liver and circulating PCSK9 is almost exclusively of hepatic origin ^6^, but the protein is expressed in several other tissues, including the intestine, kidneys, pancreas, and the nervous system ^7,8^. PCSK9 was initially cloned in cerebellar neurons undergoing apoptosis and originally called Neural Apoptosis-Regulated Convertase-1 (NARC-1) ^7^. There is evidence linking PCSK9 to lipid homeostasis as well as neuronal differentiation and inflammation in the central nervous system (CNS) ^9^. In the peripheral nervous system (PNS), PCSK9 expression has been observed in a rat Schwann cell (SC) line ^7^ and in human tibial nerves at levels deemed physiologically relevant ^10^, but the exact functions of PCSK9 in the PNS are still unknown.

Lipids play a key role in maintaining the structural and functional integrity of peripheral nerves ^11^. Despite their capacity to endogenously synthesize lipids, ^12,13^, neurons ^14^ and SC ^15^ can take up lipids from the circulation, which is underlined by the abundant expression of lipoprotein receptors in the PNS. For instance, SC express the LDLR, LRP1, and LRP8 ^16–18^. These receptors are also expressed in the dorsal root ganglia (DRG) along with the VLDLR ^19^. Given the importance of lipid homeostasis in peripheral nerves, the central role of PCSK9 in lipid metabolism, and the abundant expression of its target receptors, we hypothesized that PCSK9 could play an important role in regulating lipid metabolism in the PNS. In the present study, we show that PCSK9 is expressed in peripheral sensory neurons and in SCs. We found that PCSK9 deficiency in mice markedly reduces their behavioral responses to thermal and mechanical pain stimuli. We also observed a significant increase in CD36 protein levels and lipids content in peripheral nerves of mice lacking PCSK9. Lipid metabolism and energy production alteration were associated with mitochondrial dysfunction and axonal swelling in these mice. Together, our data demonstrate that PCSK9 plays a critical role in the regulation of lipid homeostasis in the peripheral nervous system by regulating the expression and function of CD36.

## Results

### PCSK9 is expressed in the peripheral nervous system

We assessed the expression of PCSK9 in DRG and sciatic nerves from adult wild-type (WT) mice. Using multiplex *in situ* hybridization we detected PCSK9 mRNA transcripts in both sections of DRG neurons and Schwann cells (SC) of the sciatic nerve where they respectively co-localized with the neuronal marker NeuN and the SC marker Sox10 (Figure 1A-B). The specificity of PCSK9 mRNA probes was confirmed in liver sections from WT and PCSK9 KO mice (Supplemental figure 1A). Consistently, PCSK9 protein was also detected in mouse DRG neurons expressing Neurofilament (NF) (Figure 1C) and in the sciatic nerves, where it colocalized both with the pan-SC marker S100β (Figure 1D) and the non-myelinating SC marker L1CAM (Supplemental figure 1B). These results were confirmed by analyzing PCSK9 expression in cultured primary sensory neurons isolated from P5 wild-type mice (Figure 1E) and mice and Human primary SCs culture (Figure 1F; Supplemental figure 1C). Next, we quantified PCSK9 protein levels in wild-type spinal cord, DRG, sciatic nerves, and liver extracts by ELISA, using PCSK9 KO mice as negative controls. We determined that while PCSK9 protein was not detectable in spinal cord extracts, significant levels were detected in DRG (0.46±0.11pg/µg) and sciatic nerve (0.29±0.03pg/µg) extracts, at lower concentrations than those found in the liver (0.92±0.11pg/µg) and in plasma (158±54 ng/ml) (Figure 1G). All together, these observations demonstrate that PCSK9 is endogenously expressed in sensory neurons and SC of the PNS.

**Figure 1.**
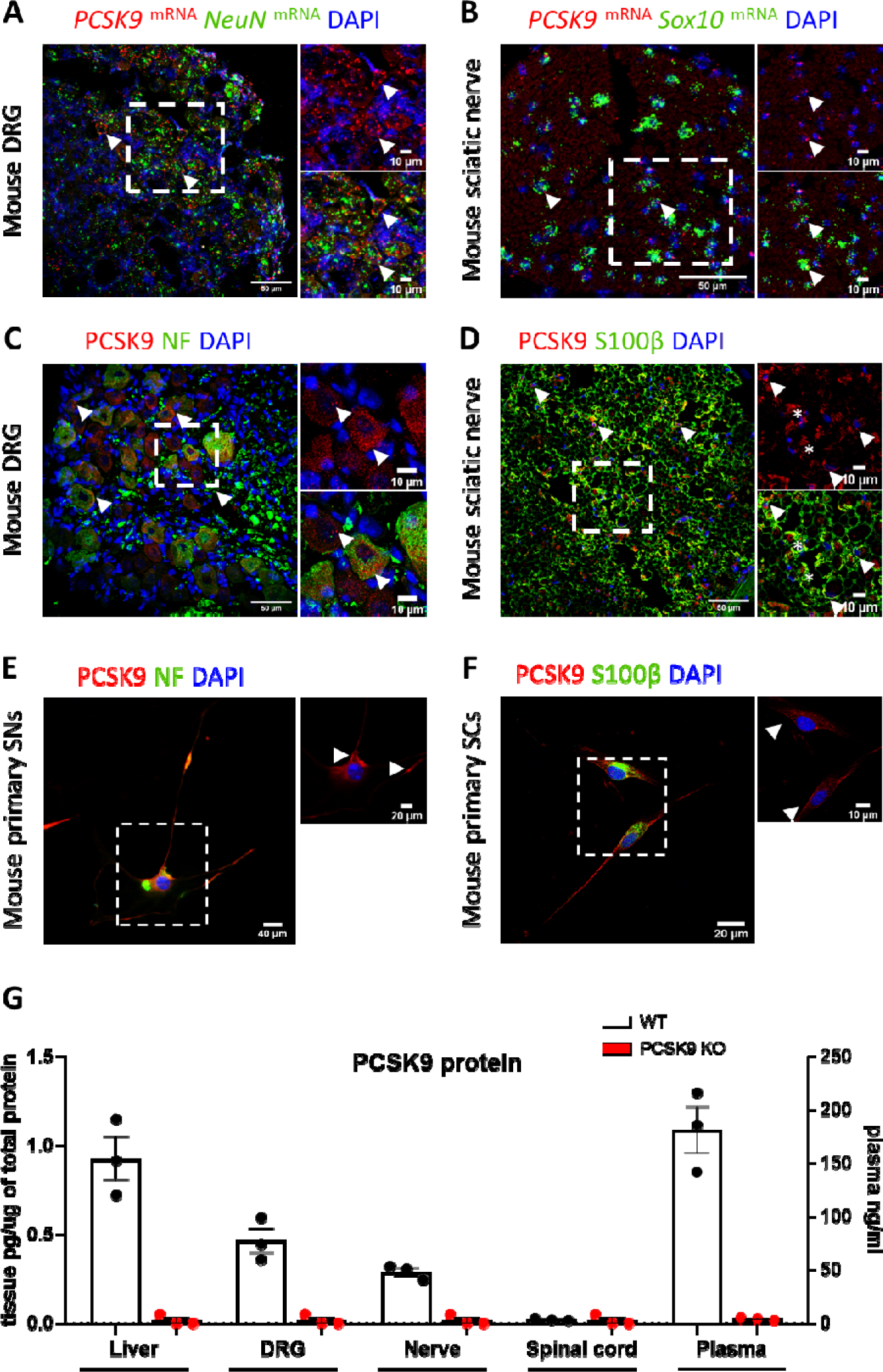
PCSK9 expression in the peripheral nervous system of mice. **(A)** PCSK9 mRNA expression in mouse dorsal root ganglia and **(B)** sciatic nerve sections. **(C)** PCSK9 protein expression in mouse dorsal root ganglia and **(D)** sciatic nerve (arrowhead: PCSK9 expression; asterisk: axonal PCSK9 expression. **(E)** PCSK9 expression in mouse primary sensory neurons and **(F)** Schwann cells (arrowhead: PCSK9 expression). **(G)** PCSK9 protein concentration in spinal cord, nerve, DRG, liver and plasma in WT mice with the corresponding negative controls from PCSK9 KO mice; *n* = 3 mice per group. Data are shown as mean ± SEM.

### PCSK9 deficiency impairs thermal and mechanical pain sensitivity

To unravel the pathophysiological consequences of PCSK9 deficiency in the functioning of the PNS, we conducted a series of behavioral analyses to comparatively assess the sensory and motor acuity of wild-type and PCSK9 knockout (PCSK9 KO) mice. At 10 weeks of age, PCSK9 KO mice exhibited reduced sensitivity to mechanical stimuli compared to wild-type animals, evidenced by an increase in paw withdrawal threshold and a decrease frequency of paw withdrawal in response to an innocuous stimulus with von Frey filaments (Figure 2A-B). We also observed a significant increase in thermal pain sensitivity in PCSK9 KO mice compared to control mice (Figure 2C). In contrast, at this stage, no clear defects were detected in their responses to noxious stimuli using either large-diameter von Frey filaments or the pinprick test (Figure 2B-D). At 24 weeks of age, in addition to the reduction of light mechanical (Figure 2E-F) and thermal pain (Figure 2G) sensations, PCSK9 KO mice also showed a significant reduction in responses to acute mechanical pain stimuli (Figure 2F-H). It is noteworthy that PCSK9 KO mice at both ages did not show any significant impairment in response to dynamic touch sensation, nor did they present with impaired motor coordination (Supplemental figure 2A-D). Taken together, these observations indicate that the absence of PCSK9 in mice impairs thermal and mechanical pain sensations.

**Figure 2.**
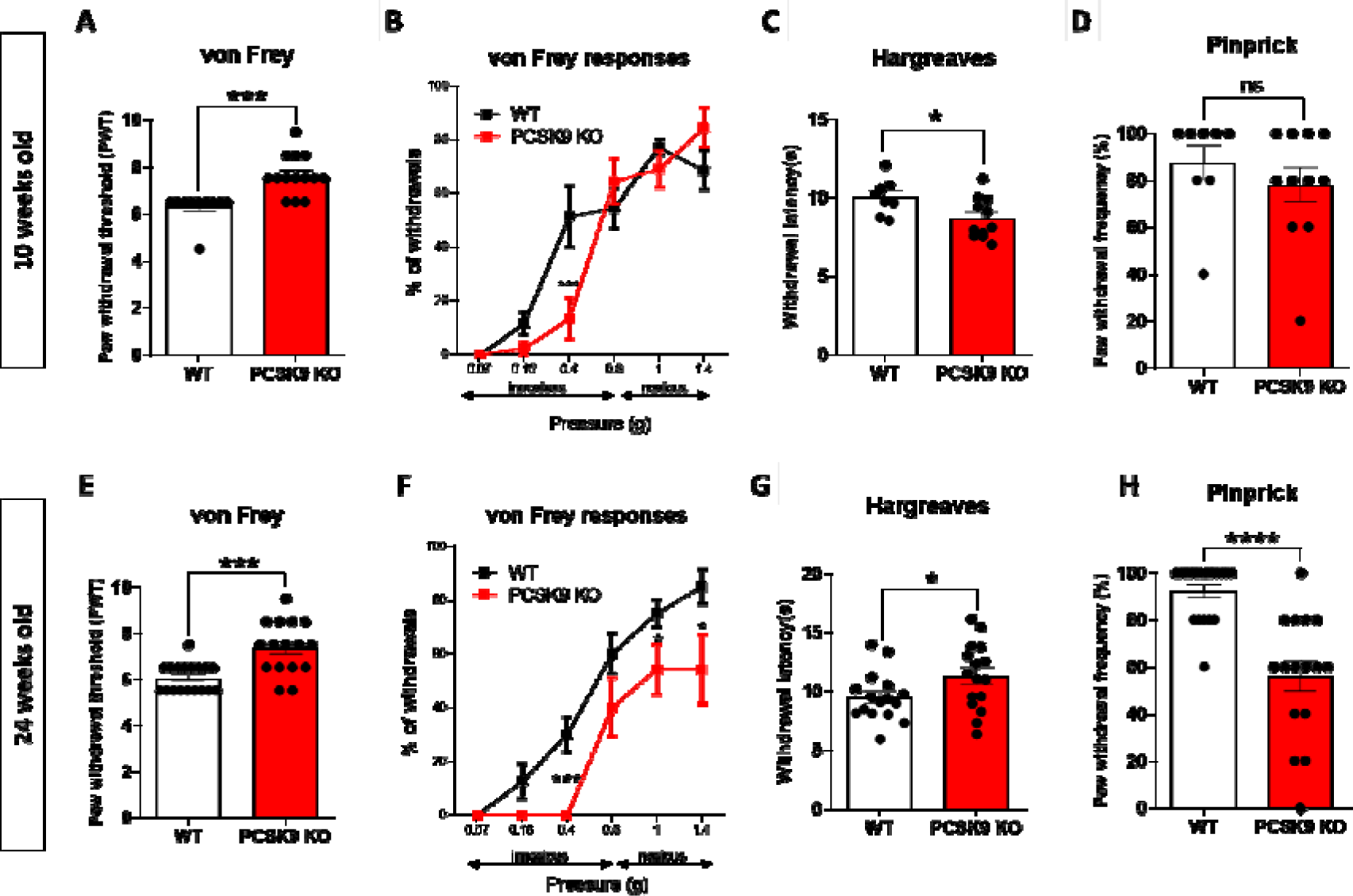
PCSK9 deficiency is associated with sensory behavioral abnormalities. **(A)** 10-week-old WT (n=10) and PCSK9 KO (n=14) mice were tested for mechanical pain sensation with the von Frey test presented as paw withdrawal threshold and **(B)** withdrawal percentage. **(C)** Thermal pain sensation in 10-week-old WT (n=8) and PCSK9 KO (n=11) mice with the Hargreaves test. **(D)** Acute mechanical pain sensation in 10-week-old WT (n=8) and PCSK9 KO (n=11) mice with the pinprick test. **(E)** 24-week-old WT (n=18) and PCSK9 KO (n=16) mice were tested for mechanical pain sensation with the von Frey test presented as paw withdrawal threshold and **(F)** withdrawal percentage. **(G)** Thermal pain sensation in 24-week-old WT (n=16) and PCSK9 KO (n=16) mice with the Hargreaves test. **(H)** 24-week-old WT (n=18) and PCSK9 KO (n=16) mice were tested for acute pain sensation with the pinprick test. Data are expressed as means ± SEM and were analyzed by unpaired t test (A, C, D, E, G and H) or 2-way ANOVA with Tukey’s multiple comparisons test (B and F). *P* > 0.05 (ns), \**P* < 0.05, \*\*\**P* < 0.001, and \*\*\*\**P* < 0.0001.

### PCSK9 deficiency alters the morphology and the conduction velocity of peripheral nerve fibers

Given that pain sensation is transmitted by small nerve fibers innervating the epidermis, we comparatively assessed intraepidermal nerve fiber density (IENFD) in the foot skin of wild-type and PCSK9 KO mice using PGP9.5 as a marker of terminal nerve fibers. Although we found similar IENFD between control and PCSK9 KO mice at 13±5 and 11±6 fibers/mm respectively, *p= 0.3903* (Figure 3A-B), we noticed a significant decrease in the number of specialized SC called nociceptive SC ^20^ (nSC) in PCSK9 knockout mice compared with wild-type, at 3±1 and 7**±**1 cells/mm, respectively, *p< 0.0001* (Figure 3C-D) by using an immunostaining for L1CAM protein that we found highly expressed in these cell types. This observation was confirmed by the quantification of nSCs expressing PGP9.5/DAPI (Supplemental figure 3A-C). We then performed electron microcopy analyses of sciatic nerve sections from these animals and we observed axonal swelling of small C-fibers at the level of Remak bundles in PCSK9 KO mice (Figure 3E). Quantitative analyses of Remak bundles revealed a shift in the proportion of these C-fibers towards an increase in axon diameter (Figure 3F). This was characterized by a significant increase in axons diameter in PCSK9 KO mice with 2.5±2% of their axons having a diameter above 1.5μm, compared with only 0.2±0.2% for wild-type mice, *p= 0.0221* (Figure 3G). Similar morphological differences between these mice were also observed in their sural nerve that mostly contain non-myelinated axons (data not shown). Remak bundles normally consist in a single non-myelinating SC that ensheaths several small C-type axon fibers separately ^21^. In PCSK9 knockouts, most axons within Remak bundles appeared to fasciculate together (Figure 3E), with an abnormal presence of vacuoles within these bundles (Supplemental figure 3D). We next comparatively assessed myelinated axons number and morphology in sciatic nerve sections from WT and PCSK9 KO mice. We did not observe any significant difference in the number of myelinated small-diameter axons (1 to 5μm) and myelinated large diameter axons (≥ *5 μm*) between WT and PCSK9 KO mice (Figure 4A).

**Figure 3.**
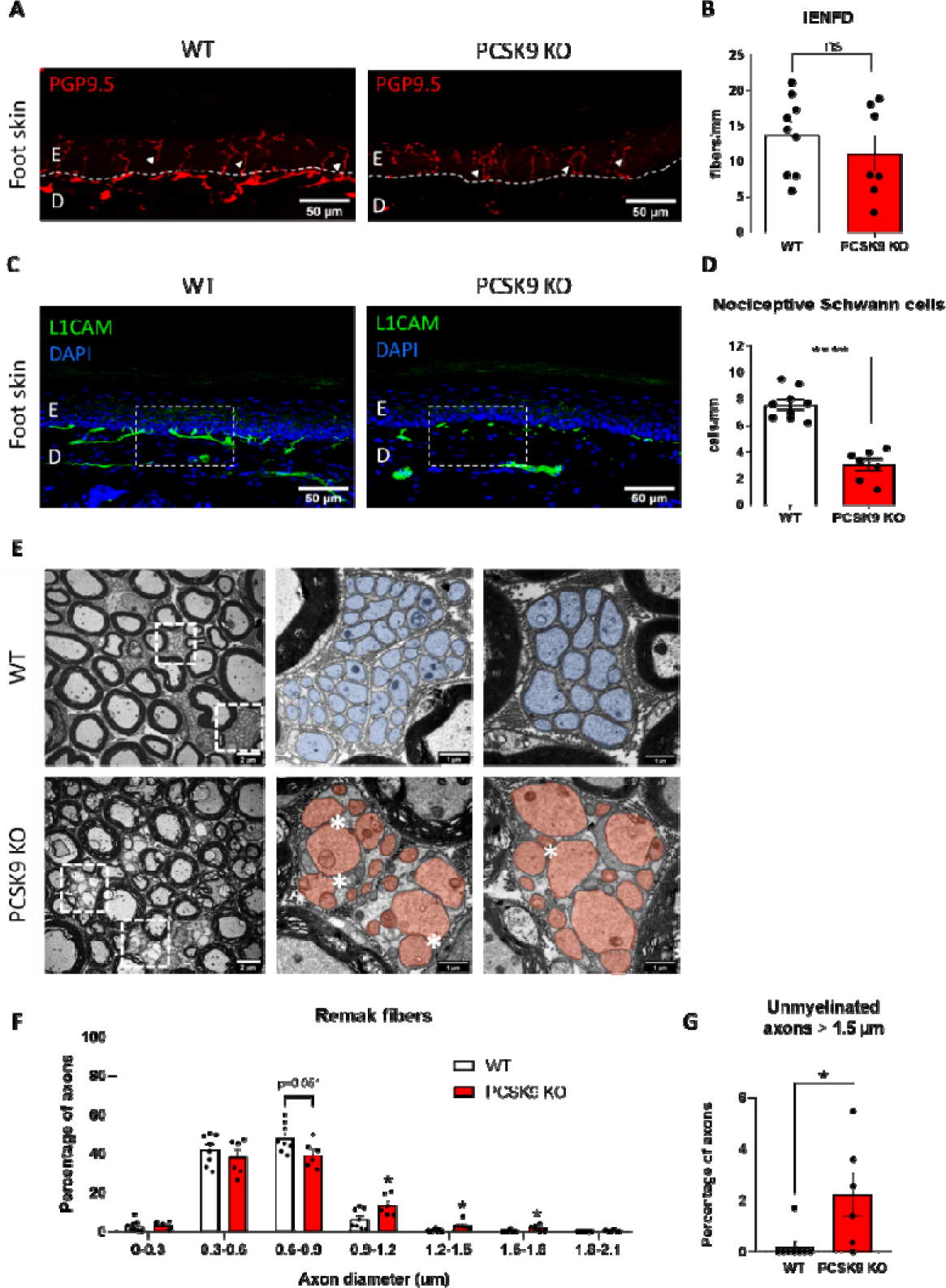
PCSK9 deficiency is associated with loss of nociceptive Schwann cells and C-fiber axonal swelling. **(A)** Representative image of intraepidermal nerve fiber density (IENFD) in the foot skin with an immunostaining against PGP9.5 (red). **(B)** Quantification of IENFD in WT (n=9) and PCSK9 KO (n=7). **(C)** Representative image of nociceptive Schwann cells in the foot skin with an immunostaining against L1CAM (green). **(D)** Quantification of numbers of nociceptive Schwann cells between WT (n=9) and PCSK9 KO (n=7) mice. **(E)** Representative photomicrographs of Remak bundle structure by transmission electron microscopy in WT (small C fibers are highlighted in blue) and PCSK9 KO mice (small C fibers are highlighted in pink); (*) indicates axon-axon contacts. **(F)** Distribution of axon diameter in Remak bundles between WT (n=8) and PCSK9 KO (n=6) mice. **(G)** Percentage of small unmyelinated C-fibers with a diameter greater than 1.5 µm in WT (n=8, ≥2035 axons were analyzed) vs. PCSK9 KO (n=6, ≥1727 axons were analyzed). Data are represented as mean ± SEM and analyzed by unpaired t test (B, D, and G) and Multiple t test (F). *P* > 0.05 (ns), \**P* < 0.05 and \*\*\*\**P* < 0.0001.

**Figure 4.**
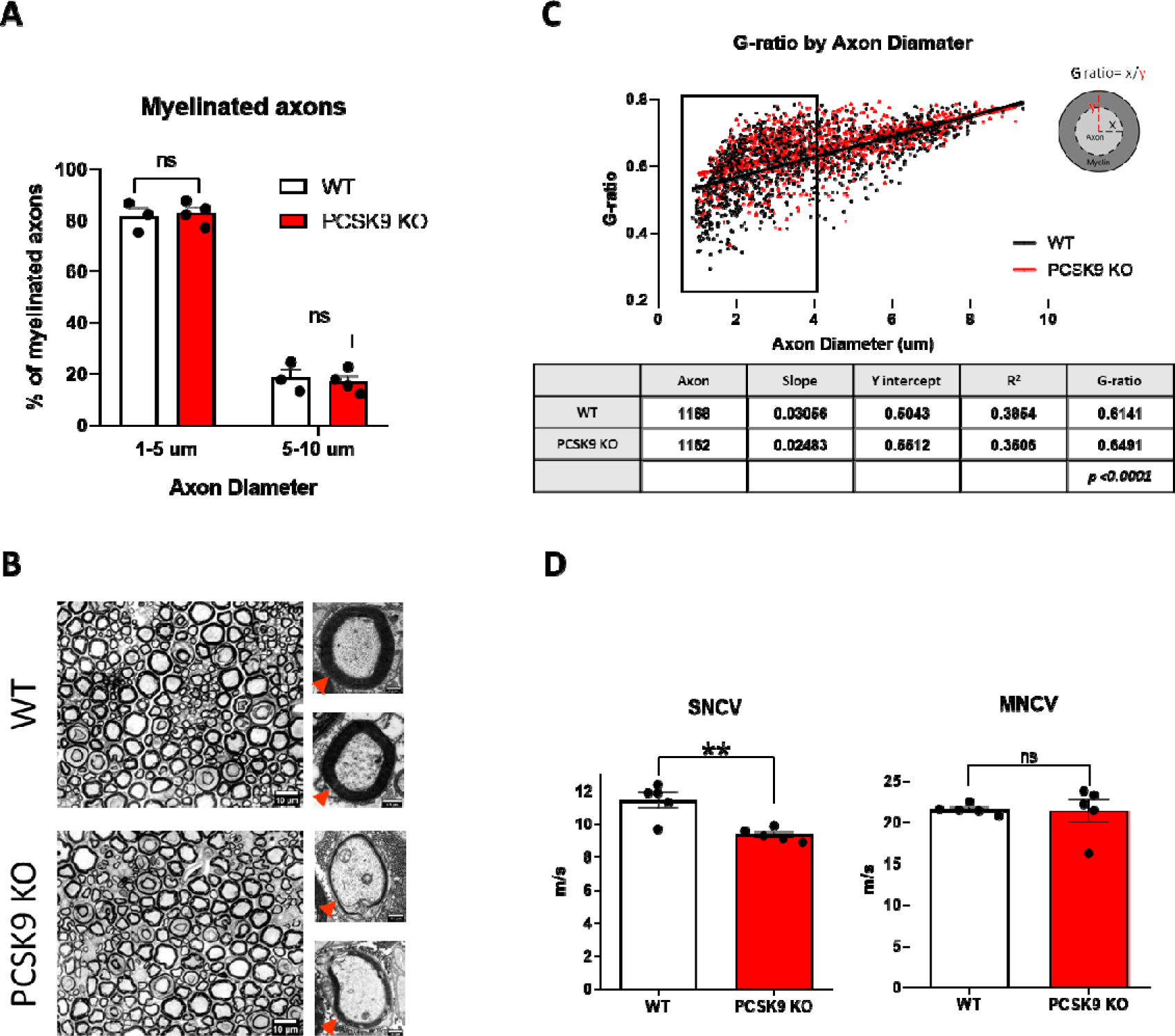
PCSK9 deficiency is associated with structural and physiological changes in myelinated A fibers. **(A)** Proportions of myelinated axons distributed according to axon diameter in both WT (n=3) and PCSK9 KO (n=4) mice. **(B)** Representative images of toluidine blue staining of myelin in WT and PCSK9 KO mice (red arrowheads indicates myelin). **(C)** G-ratio analysis with a summary table in WT (n=4, ≥1168 axons were analyzed) vs. in PCSK9 KO (n=3, ≥1162 axons were analyzed). **(D)** Sensory nerve conduction velocity (SNCV) and motor nerve conduction velocity (MNCV) compared between WT (n=5) and PCSK9 KO (n=5) mice. Data are represented as mean ± SEM and statistically analyzed by Multiple t test (A and C) and unpaired t test (D). *P* > 0.05 (ns), \*\**P* < 0.01.

Interestingly, our analysis of the myelination status of these fibers revealed that small myelinated A-fibers (≤ *5 μm)* in PCSK9 KO mice had a significant increase in their G-ratio reflecting a decrease in myelin thickness, compared with wild-type mice (Figure 4B-C). Importantly, these morphological differences observed in PCSK9 KO mice translate at the functional level into a significantly reduced sensory nerve conduction velocity (9.36 ± 0.39 vs. 11.45 ± 1.04 m/s, *p=0.0031*) but similar motor nerve conduction velocity (21.38 ± 3.04 vs. 21.56 ± 0.61 m/s, *p=0.9*), compared with wild-type mice, respectively (Figure 4D). Altogether, these data show that although PCSK9 knockout mice did not exhibit any change in the density of terminal nerve fibers in the skin, the number of nociceptive SCs was significantly reduced. Moreover, while the peripheral nerves of mutant mice contain normal numbers of axons, they exhibit axonal swelling of C-fibers present in the Remak bundles, and a decrease in myelin thickness of small A-fibers paralleling the reduction of sensory nerve conduction velocity. Thus, PCSK9 KO mice present with alterations in peripheral nerve morphology and conduction properties.

### PCSK9 modulates CD36 but not LDLR expression in peripheral nerves

To gain insights into the molecular mechanism(s) underlying the sensory and morphological phenotypes observed in PCSK9 knockout mice, we assessed the expression of the main PCSK9 targets in the sciatic nerves of these animals. Surprisingly, we did not detect any change of the level of LDLR in the peripheral nerves of PCSK9 knockout mice compared to WT animals (Figure 5A-D), in contrast to what was observed in the liver (Supplemental figure 4A, B). Similar results were obtained by analyzing the expression of VLDLR, LRP1 and apoER2 receptors (Supplemental figure 5A-C). In contrast, we strikingly observed a significant increase in the expression of CD36 in sciatic nerves of PCSK9 KO compared with wild-types (Figure 5E-H), as in the liver (Supplemental figure 4A, C). We established that up-regulation of CD36 in the peripheral sciatic nerve of PCSK9 knockout mice was prominent in nerve fibers as well as myelinating and non-myelinating SC (Supplemental figure 6A-B). Of note, in the DRG, CD36 was expressed at a very low levels, and its expression as well as that of the LDLR were not significantly altered in PCSK9 KO mice (Supplemental figure 7A-D). Our findings thus demonstrate that the absence of PCSK9 specifically induces the upregulation of CD36 but not that of the LDLR or other target receptors in peripheral nerves.

**Figure 5.**
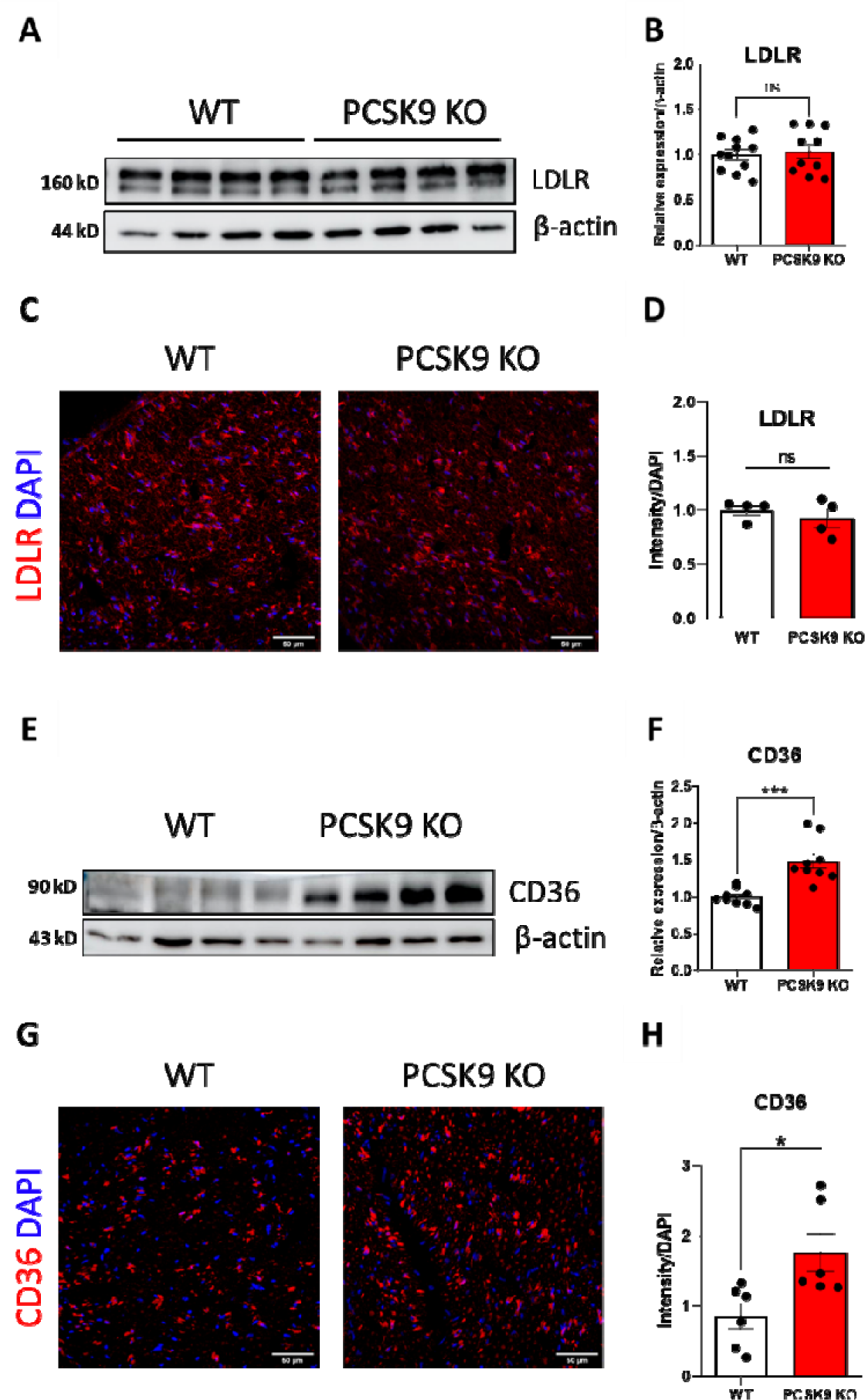
PCSK9 deficiency results in increased of CD36 expression in peripheral nerves. **(A)** LDLR expression in the sciatic nerve by immunoblotting. **(B)** Quantification of LDLR expression in 24-week-old WT (n=11) and PCSK9 KO (n=10). **(C)** Immunohistochemistry for LDLR in sciatic nerve sections of 24-week-old WT and PCSK9 KO mice. **(D)** Immunohistochemistry quantification of LDLR expression in 24-week-old WT (n=4) and PCSK9 KO (n=4) nerve sections. **(E)** CD36 expression in the sciatic nerve by immunoblotting. **(F)** Quantification of CD36 expression in 24-week-old WT (n=9) and PCSK9 KO (n=9). **(G)** Immunohistochemistry for CD36 in sciatic nerve sections of 24-week-old WT and PCSK9 KO mice. **(H)** Immunohistochemistry quantification of CD36 expression in 24-week-old WT (n=6) and PCSK9 KO (n=6) nerve sections. Data are represented as mean ± SEM and statistically analyzed by unpaired t test. *P* > 0.05 (ns), \**P* < 0.05 and \*\*\**P* < 0.001.

### PCSK9 deficiency is associated with lipid accumulation in peripheral nerves

We next evaluated whether the upregulation of the fatty acid transporter CD36 in peripheral nerves altered their lipid contents. We measured cholesterol and triglyceride concentrations in sciatic nerve extracts and observed an increase of 30% and 50% respectively, in PCSK9 knockout comparatively to wild-type mice (Figure 6A-B). Liver lipid contents were also measured and were consistent with the results of previous studies ^22^ (Supplemental figure 4D-E). In contrast, we did not observe any change in cholesterol and triglyceride concentration in DRG extracts from control and PCSK9 KO mice (Supplemental figure 7E-F). In peripheral nerves, cholesterol and triglyceride contents remarkably correlated with CD36 protein expression (Supplemental figure 8A-B). Furthermore, neutral lipid staining of sciatic nerve sections showed an accumulation of lipid droplets in PCSK9 knockout compared to control animals (Figure 6C-D). Structurally, by electron microscopy analysis we observed that PCSK9 KO peripheral nerves displayed lipid droplets within Remak bundles (Figure 6E). Taken together, these results indicate that PCSK9 deficiency leads to an abnormal lipid accumulation within peripheral nerves.

**Figure 6.**
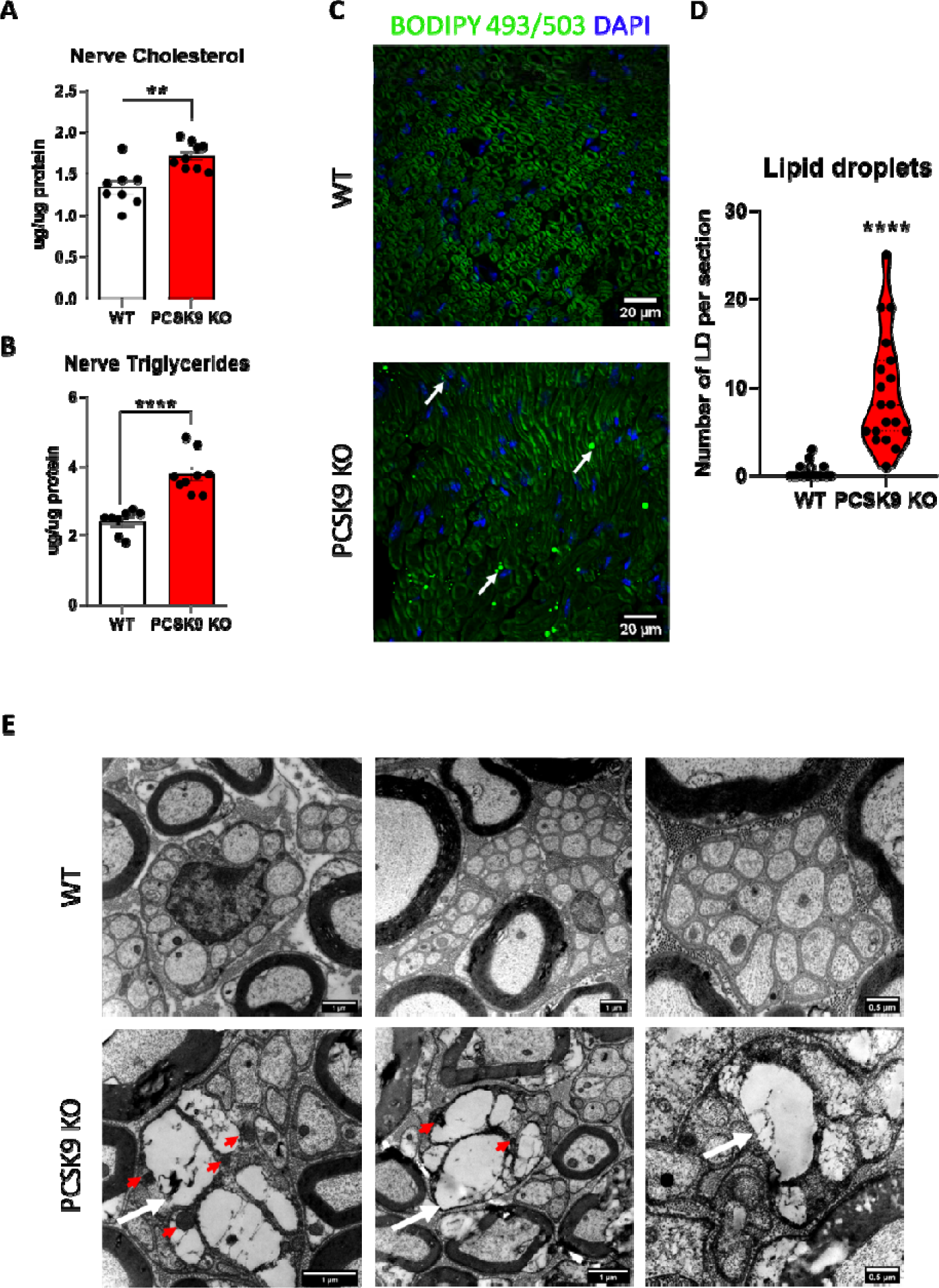
PCSK9 deficiency results in increased lipids concentration in peripheral nerves. **(A)** Cholesterol levels in the sciatic nerve of 24-week-old WT (n=8) and PCSK9 KO (n=9) mice. **(*B*)** Triglyceride levels in the sciatic nerve of 24-week-old WT (n=8) and PCSK9 KO (n=9) mice. **(C)** Visualization of lipid droplets in sciatic nerve sections using BODIPY 493/503 staining (arrows: lipid droplets). **(D)** Quantification of the number of lipid droplets per section from 24-week-old WT (n=3) and PCSK9 KO (n=4) mice. **(E)** Representative photomicrographs of sciatic nerve sections by transmission electron microscopy in WT and PCSK9 KO mice are shown. White arrows indicate lipid droplets; red arrows show the associated mitochondria. Data are represented as mean ± SEM and statistically analyzed by unpaired t test. \*\**P* < 0.01 and **** *P* <0.0001.

### Mitochondrial defects in peripheral nerves of PCSK9 KO mice

To fully unravel the pathophysiological onset of the sensory phenotype of PCSK9 knockout mice, we next performed an untargeted proteomic investigation of their sciatic nerves. We observed that PCSK9 knockout mice showed a significant upregulation of several proteins involved in lipid mitochondrial metabolism and a downregulation of cytoskeletal proteins compared to wild-type mice (Figure 7A). Major upregulated proteins are implicated in β-oxidation such as Carnitine O acetyl transferase (Crat) and Trifunctional enzyme subunit alpha (Hadha), enzymes in the TCA cycle like Isocitrate dehydrogenase [NAD] subunit alpha (Idh3a), and proteins of the Electron Transport Chain including Complex I (Ndufs5), Complex III (Uqcrc2, Uqcrb) and Complex IV (Cox5b). Interestingly, Caveolin-1, a protein necessary for the localization and functioning of CD36 at the plasma membrane, was also upregulated in PCSK9 KO mice, in line with CD36 expression results. The proteomic analysis also revealed a downregulation of proteins involved in proteasomal catabolic processes such as Psma1, Psmb2, and Psma5, as well as microtubule organization proteins including Map4, Mapt, Msn, and Sorbs3 as well as the myelin P2 protein (PMP2) constitutive of peripheral nerve myelin (Figure 7A). Biological processes enrichment analysis of PCSK9 knockout nerves proteome indicated increased overall processes of lipid metabolism and electron transport chain processes and downregulation of processes involved in microtubule cytoskeleton organization and the capacity to respond to oxidative stress (Figure 7B) and most of the identified upregulated proteins are clustered as mitochondrial components (Supplemental figure 8C). To corroborate these results, we analyzed mitochondria number and morphology in section of sciatic nerve by electron microscopy. Interestingly, we observed an increase number of mitochondria in Remak bundle nerve fibers of PCSK9 KO mice compared to control (Figure 7C-D) and an increased proportion of abnormal mitochondria, which is representative of mitophagic processes (Figure 7D-E). Taken together, these observations indicate that PCSK9 knockout mice present with structural mitochondrial defects in the peripheral nerves.

**Figure 7.**
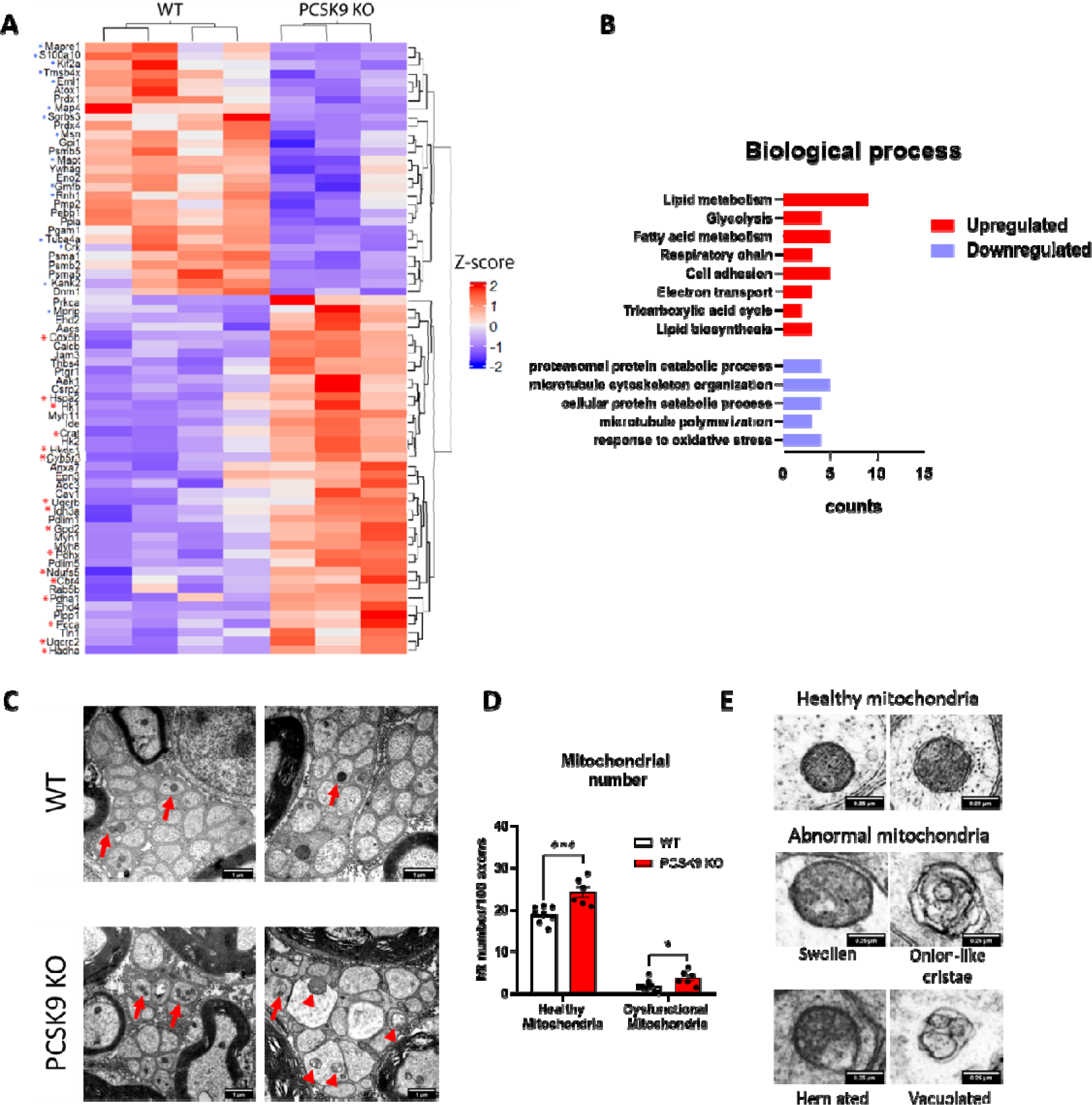
Proteomic profile and mitochondrial abnormalities of PCSK9 KO nerves. **(A)** Heatmap showing relative protein expression values (*z*-score) of *n*=62 proteins that are differentially expressed in the nerves of WT (n=4) compared to PCSK9 KO (n=3) mice (red asterisk represents upregulated mitochondrial proteins, blue asterisk represents downregulated microtubule proteins). **(B)** Gene ontology annotation of the top upregulated (red) and downregulated terms in biological processes sorted by P-value. **(C)** Representative TEM images of Remak bundles (red arrows: healthy mitochondria, red arrowheads: abnormal mitochondria). **(D)** Quantification of total mitochondrial numbers with the ratio of abnormal mitochondria per 100 axons in WT (n=8, ≥2035 axons were analyzed) and PCSK9 KO (n=6, ≥1727 axons were analyzed). **(E)** Representative images of healthy mitochondria and abnormal mitochondria. Data are represented as mean ± SEM and statistically analyzed by unpaired t test. \**P<*0.05, \*\*\**P*≤0.001.

## Discussion

In this study, we described the expression of PCSK9 in the peripheral nervous system at the level of sensory neurons and Schwann cells. We found that PCSK9 gene deletion induces a neuropathic phenotype characterized by a reduction in thermal and mechanical pain sensations at 24 weeks old mice. These sensory deficits were associated with a reduction in sensory nerve conduction properties without significant alteration of the number of sensory axons in the nerve nor in the density of their terminal endings in the skin. In contrast there was a reduction in the number of terminal nociceptive SCs at the border between the dermis and the epidermis. We established that PCSK9 gene deletion induces an upregulation of the fatty acid transporter CD36 associated with an abnormal lipid’s accumulation in the peripheral nerves. We also showed by proteomic and histological analysis that PCSK9 deficiency leads to structural mitochondrial defects. In light of these observations, we propose a model in which PCSK9 through the regulation of CD36 limits the internalization of fatty acid and accumulation of lipid storage in the peripheral nerves (Graphical abstract).

Here we show for the first time that PCSK9 is physiologically expressed in the PNS at the level of the sensory neurons and in both myelinating and non-myelinating SCs, which had initially been suggested by the detection of PCSK9 mRNA in a rat SC line ^7^ and in human tibial nerves ^10^. We also observed that mice lacking PCSK9 show sensory defects characterized by a decrease in their ability to perceive light mechanical pain and an increase in thermal pain sensitivity at 10 weeks of age, and subsequently become less sensitive to acute mechanical and thermal pain stimuli at 24 weeks.

In line with their neuropathic phenotype, we observed structural alterations at the level of terminal endings of the peripheral nerves in the skin of PCSK9 knockout mice. Intriguingly, while we did not observe any significant decrease in IENFD in the skin, we noticed a significant reduction in the number of nociceptive Schwann cells, a non-neuronal cutaneous cell subtype in close relationship with sensory endings known to play a critical role in tactile sensations ^20^. At the level of sciatic nerves, we observed axonal swelling of unmyelinated C-fibers in Remak bundles, as well as a reduction in the thickness of the myelin sheath around small diameter A-myelinated axons in PCSK9 KO mice. These two types of fibers, referred to as unmyelinated C-fibers and myelinated Aδ-fibers, respectively, are specifically involved in the transmission of mechanical pain ^23,24^. The combination of these structural abnormalities, including the loss of nociceptive Schwann cells, the swelling of small C fiber axons, and the hypomyelination of A-δ fibers, appears to be at the origin of the sensory deficits observed in PCSK9 KO mice.

We also showed that PCSK9 knockout mice present an increased expression of CD36 in peripheral nerve SCs and axons, whereas the expression of the LDLR, VLDLR, ApoER2 and LRP1 were unaltered in these tissues comparatively to wild-type mice. The absence of modulation of LDLR abundance in PCSK9 knockout mice peripheral nerves is surprising, given the overt role that PCSK9 plays in LDLR intracellular degradation in hepatocytes and many other cell types ^25,26^. Nevertheless, the lack of LDLR modulation in the brain and cerebellum of PCSK9 KO mice has previously been reported by others ^27–29^. Although the mechanism is still unclear, several proteins have been proposed as intermediates to either enhance or impede PCSK9-mediated degradation of LDLR. Indeed, Annexin 2 ^30,31^ and glypican-3 ^32^ were shown to inhibit PCSK9 binding to LDLR, whereas cyclase-associated protein 1 ^33^ enhances its binding capacity. The differential expression of these proteins between the hepatic and nervous system could potentially explain the distinct regulatory functions of PCSK9 on LDLR between these tissues. In contrast, the increase in CD36 expression has been reported in many organs of PCSK9 knockout mice, including the liver, adipose tissue, heart, and kidneys ^34,22,35,36^. The increase in CD36 expression resulting from PCSK9 gene deletion in these tissues was systematically associated with an abnormal accumulation of lipid droplets^34,22,35,36^. Accordingly, we also found an accumulation of lipids in the peripheral nerves of PCSK9 knockout mice proportional to the levels of CD36 expression. Of note, the deteriorating sensory phenotype from 10 to 24 weeks discussed earlier could potentially be attributed to an increase in toxicity caused by the gradual accumulation of lipids over time. Collectively, these observations support a role for PCSK9 in controlling CD36 levels in peripheral nerves, thereby protecting nerve cells against excessive lipid uptake and accumulation that can lead to neuronal lipotoxicity.

In line with this hypothesis, we found that PCSK9 knockout mice nerves present with an increased expression of mitochondrial enzymes involved in β-oxidation, enzymes of the TCA cycle, as well as proteins of the electron transport chain. CD36 induced accumulation of lipids in peripheral nerves may indeed overwhelm the capacity of SCs and axons to metabolize these compounds through mitochondrial β-oxidation, the TCA cycle and oxidative phosphorylation to generate ATP. We also found that proteins involved in proteasomal catabolic processes and microtubule biology were downregulated, both parameters being indicative of neurodegenerative processes ^37,38^. Furthermore, we found structurally abnormal and dysfunctional mitochondria within unmyelinated Remak bundle fibers of PCSK9 deficient mice. In line with these observations, an increased proportion of dense depolarized mitochondria in peripheral axons has also been reported in mouse models of diabetic neuropathy ^39,40^ and cardiac mitochondrial metabolism was found altered in cardiomyocytes of PCSK9 knockout mice, in addition to enhanced CD36 expression and the presence of lipid droplets in the heart of these animals ^35^. These observations show that in the absence of PCSK9, peripheral nerves exhibit increased CD36 expression and an abnormal accumulation of lipids. We propose that this excess of lipids leads to mitochondrial dysfunction in small nerve fibers, and can contribute to the sensory neuropathic phenotype observed in these animals.

Interestingly, these results contrast with the fact that over the last decade only one documented case of peripheral neuropathy has been to be linked to PCSK9 inhibitors usage ^41^. Long term trials conducted with evolocumab and alirocumab, two PCSK9 inhibitor antibodies have not demonstrated any link to negative neurocognitive events ^4,42^. However, there was no specific evaluation of sensory motor endpoints. Of note, these drugs specifically antagonize the domain of interaction between PCSK9 and the LDLR but not with the fatty acid transporter CD36 ^1,36^, and theoretically would thus not alter the regulatory effect of PCSK9 on CD36. In addition, these drugs as well as the novel anti-PCSK9 siRNA inclisiran only target circulating PCSK9 ^6,43^. Given that PCSK9 KO mice totally lack PCSK9 in the circulation as well as in all tissues, we cannot rule out that the neuropathic phenotype of PCSK9 knockout mice might stem from the absence of local PCSK9 within SCs and/or sensory neurons rather than from the absence of circulating PCSK9.

Nevertheless, the discovery of a role for PCSK9 in peripheral nerve lipid homeostasis, and its modulation through the CD36 regulation open news avenues of research in certain cases of acquired demyelinating disorders of peripheral nerves (i.e. Guillain-Barre Syndrome, Anti-Myelin Associated Glycoprotein Neuropathy) as well as in nerve injury and remyelination, where lipids are considered limiting resources ^44,45^. Future investigations are necessary to better understand the underlying mechanisms of modulation of PCSK9 and of its target receptors in such pathologies in order to design news targeted therapeutical approaches for these diseases.

## Methods

### Sex as a biological variable

Our study exclusively examined male mice with the rationale that sex-specific phenotypes in mice lacking PCSK9 have been observed previously ^46,47^. It is unknown whether the findings are relevant for female mice.

### Mice

*B6;129S6-Pcsk9tm1Jdh/J* (*Pcsk9* KO) (Jackson Laboratory, Bar Harbor, ME) and *C57BL/6J* wild type (WT) mice were kept under a controlled light/dark cycle (12 hours of light/12 hours of dark), under temperature-controlled conditions (21°C) and standard humidity. Mice were fed chow diet ad libitum with free access to water. Mice were anesthetized with isofluorane and euthanized by intracardiac puncture. Blood samples were collected for plasma isolation by centrifugation and tissues (liver, DRG, sciatic nerves, foot pads) were snap frozen in liquid nitrogen. Plasma and tissue samples were stored at −80°C until use. All mice experiments were performed at 24 weeks of age unless mentioned otherwise.

### Immunohistochemistry

Right after euthanasia, mice were perfused with 20 mL 1X PBS followed by 15 mL of 4% paraformaldehyde (PFA) dissolved in 1X PBS. Sciatic nerves, foot pads, and DRGs were dissected and fixed overnight in 4% PFA at 4°C. Tissues samples were washed 3 times (10 min) in cold PBS, cryoprotected overnight in a PBS solution containing 30% sucrose, and embedded in OCT compound, and stored at −80 °C, until use. Frozen tissues were cut into 14-μm sections (20-μm for foot pads), placed onto glass slides, dried on a heating plate for 10 mins. The slides were stored at −20°C until use. Sections were rehydrated in PBS, permeabilized with PBT (PBS + 0.1% Triton X-100), blocked in donkey serum (Sigma-Aldrich, Saint-Quentin-Fallavier,France), prior to an overnight incubation with primary antibodies (Table 1). For PCSK9 immunostaining, we added an antigen retrieval step by incubating the sections for 10 min in 10 mM citrate buffer, pH 6.0 at 95°C. Slides were washed 3 times in PBT for 10 mins, incubated with fluorescent secondary antibodies (Table 1) for 1 hour, and washed again as above. Nuclei were counterstained with DAPI (0.5 µg/ml), slides were washed again and then mounted and kept at 4 °C.

**Table 1:** List of all antibodies used (IF: immunofluorescence, WB: western blot)

| Target antigen | Vendor or source | Catalog # | Experiments |
| --- | --- | --- | --- |
| Rabbit anti human pro-PCSK9 N-terminal domain | Abcam | ab135647 | IF |
| Rabbit anti-mouse PCSK9 | Cayman | 10240 | IF |
| Mouse anti-S100β | Sigma-Aldrich | S2532 | IF |
| Rat anti- L1CAM | Merck milipore | MAB5272 | IF |
| Goat anti-LDLR | R&D | AF2255 | IF, WB |
| Mouse anti-ApoER2 | Abnova | H00007804-M01 | IF, WB |
| Rabbit anti-LRP1 | Abcam | ab92544 | IF, WB |
| Goat anti-VLDLR | R&D | AF2258 | IF, WB |
| Chicken anti-NF | Merck milipore | AB5539 | IF |
| Rabbit anti-β-actin | Abcam | ab8227 | WB |
| Rabbit anti-PGP9.5 | Merck milipore | AB1761 | IF |
| Rabbit anti-CD36 | Abcam | ab124515 | IF, WB |
| Donkey anti-Rabbit Alexa 594 | ThermoFisher | A-21207 | IF |
| Donkey anti-Mouse Alexa 594 | ThermoFisher | A-21203 | IF |
| Donkey anti-Rat Alexa 488 | ThermoFisher | A-21208 | IF |
| Donkey anti-Goat Alexa 594 | ThermoFisher | A-11058 | IF |
| Donkey anti-Chicken Alexa 488 | Invitrogen | A78948 | IF |
| Donkey Anti-Goat IgG H&L (HRP) | Abcam | ab205723 | WB |
| Peroxidase AffiniPure™ Donkey Anti-Rabbit IgG (H+L) | Jackson ImmunoResearch | 711-035-152 | WB |
| Peroxidase AffiniPure™ Donkey Anti-Mouse IgG (H+L) | Jackson ImmunoResearch | 715-035-150 | WB |

### Rnascope

RNA in situ hybridization was performed using the RNAscope Multiplex Fluorescent Reagent Kit Assay (Advanced Cell Diagnostics, Inc., Newark, CA) according to the manufacturer’s instructions. RNAscope probes used were as follows: Pcsk9 (Cat No. 498111), Sox10 (Cat No. 435931), Rbfox3 (NeuN) Cat No. 313311-C2 and Ldlr (Cat No. 443701). Nuclei were ultimately counterstained with DAPI.

### Image acquisition and analysis

Images were acquired on a laser scanning confocal microscope Eclipse Ti2 (Nikon) with identical parameters (laser power, iris diameter, and gain) to acquire images from all sections.

### Mouse and human primary Schwann cell culture

Primary Schwann cell cultures were isolated as follows: Sciatic nerves of 6 mouse pups between postnatal days 2 and 4 (P2 and P4) were collected in L15 medium (Thermofisher, Massachusetts, USA) and incubated for 30 min in 2 mg/mL collagenase II and then 10 min in 0.25 % trypsin containing EDTA at 37 °C. Nerves were dissociated into single-cell suspensions with a 1mL pipet and then through an 18-G/21-G needle. Schwann cells were resuspended in primary Schwann cell medium (ScienCell #1701, California, USA) supplemented with 1 μM insulin, 2 μM forskolin, and 10ng/mL Nrg1ß ^48^.

Human primary Schwann cells isolated from human spinal nerve were commercially obtained from Sciencell (California, USA) Catalog No. 1700. Cells were maintained in primary Schwann cell medium.

### Mouse primary sensory neurons culture

Primary sensory neuron cultures were isolated as follows: dorsal root ganglia of 6 mouse pups between postnatal days 2 and 4 (P2 and P4) were collected in PBS-Glucose (0.3 µM) and incubated for 30 min in 0.125% collagenase D (Roche, Switzerland) and then 10 min in 0.25 % trypsin at 37 °C. Ganglia were dissociated into single-cell suspensions with a glass Pasteur pipette. Sensory neurons cells were resuspended in Neurobasal (NB) Gibco™21103049 (Thermofisher, Massachusetts, USA) supplemented with B27 Gibco™ 17504044 (Thermofisher, Massachusetts, USA).

### Biochemistry and western blot analyses

Cholesterol and triglyceride contents in plasma and tissue samples were determined using colorimetric assays (Diasys, Holzheim, Germany). Concentration of PCSK9 in the plasma and tissue samples were assessed by ELISA (Quantikine MPC900, R&D Systems, Minneapolis, MN). Protein extracts were obtained after lysis of tissues in 350-μL Tris HCl buffer pH=9.5 (10 mM) containing EDTA (1mM), NaCl (150 mM) supplemented with the Halt™ Protease and Phosphatase Inhibitor Cocktail (Thermofisher, Massachusetts, USA). Proteins (≈20µg) were separated by sodium dodecyl sulfate 7.5% polyacrylamide gel electrophoresis and transferred onto a nitrocellulose membrane. Membranes were blocked in tris-buffer saline containing 5% (w/v) non-fat dry milk, 0.1% (v/v) Tween 20 (TBS) for 1 h at room temperature before incubation with the primary antibodies (Table 1) overnight at 4 °C. Membranes were washed in TBS-T and incubated for 90 min at 25°C with HRP-conjugated secondary antibodies (Table 1). Immunoreactive bands were detected by chemiluminescence (Amersham ECL Prime Western Blotting Detection Reagent, GE Healthcare Life Sciences, Vélizy-Villacoublay, France) and images were acquired with an Amersham Imager 600 (GE Healthcare, Vélizy-Villacoublay, France) and analyzed with the ImageJ software (Maryland, USA). Throughout, results were normalized to β-actin expression.

### Behavioral assessment of peripheral neuropathy

#### von Frey test

Mice were placed in a plastic chamber (10 x 10 x 14 cm) on an elevated wire grid. They were habituated to the testing apparatus for 1h/day during a week before behavioral testing. The plantar surface (glabrous) of the hind paw was stimulated with a set of calibrated von Frey filaments (0.008 - 6 g) (Bioseb - *In Vivo* Research Instruments, Vitrolles, France). The paw withdrawal threshold (PWT) was determined as described previously ^49^. Von Frey responses were determined as the percentage of withdrawal using specific filament (0.07, 0.16, 0.4, 0.6, 1,1.4 g). Each filament was tested five times in increasing order.

#### Pinprick test

Mice were placed in a plastic chamber on an elevated wire grid. The plantar surface of the hind paw was stimulated with a pin gently applied without moving the paw or penetrating the skin. Pin stimulation was repeated 5 times on different areas with a 1–2 min interval between each stimulation. The percentage of positive attempts (paw withdrawal) was calculated.

#### Hargreaves test

In the Hargreaves test, we measured the response to radiant heat. Mice were placed in a plastic chamber on a glass floor and the infrared heat stimulus from the plantar test (Ugo Basile, France) was applied to the hind paw. The latency for the animal to withdraw the hind paw was measured. The beam intensity was adjusted at 20% so that control mice displayed a latency of 8–12 s. A cutoff time of 30 seconds was used to avoid tissue damage.

#### Brush test

To assess sensitivity to dynamic light touch, we used a brush test. The plantar surface of the hind paw was stimulated with a soft paintbrush by gently stroking from heel to toe. The brush stimulation was repeated 5 times with an interval of 1 min and the percentage of paw withdrawals from 5 stimuli was calculated.

#### Rotarod test

Gross motor ability and coordination were assessed using a rotarod apparatus (Bioseb-Vitrolles, France). Mice were first placed on the apparatus for 30 s with no rotation and thereafter for 2 min at a constant low-speed rotation (4 rpm). Those that fell from the rod at 4 rpm were placed again on it until they were able to stay for 1 min. Mice were tested in accelerating conditions from 4 to 40 rpm over 2 min period. The time during which the animal walked on the rod before falling was collected (maximum value: 120 s). The occurrence of two consecutive passive rotations, i.e., without walking but accompanying the rod, was considered as a fall. Each mouse was tested three times per day with inter-trial intervals of 5 min.

### Nerve conduction studies (NCV)

Mice (24-week-old) were anesthetized with isoflurane at doses of 4–5% for induction and 1– 2% for maintenance. The core temperature was maintained at 34 °C with a heating pad. Stainless steel needle electrodes (Natus Biomedical, Middleton, Wisconsin) were cleaned with 70% alcohol between animals. Using the Natus UltraPro S100 EMG/NCS system and the Keypoint 4 software, sensory NCV was determined by dorsal paw recordings and stimulating electrodes on the ankle. For sensory NCV, the latency of onset (milliseconds) of the sensory nerve action potential after supramaximal antidromic stimulation of the sural nerve at the ankle was divided by the distance between the recording and stimulation electrodes (measured in millimeters using a Vernier caliper). Motor NCV was calculated by subtracting the distal from the proximal latency (measured in milliseconds) from the stimulus artifact of the take-off of the evoked potential, and the difference was divided by the distance between the two stimulating electrodes (measured in millimeters using a Vernier caliper).

### Intra epidermal nerve fiber density (IENFD) and Nociceptive SC quantification

IENFD quantification was performed on WT and PCSK9 KO mice. Both right and left plantar surface of the hind paw were collected, fixed overnight with 4% PFA. Tissues were rinsed with 1X PBS and then incubated in 30% sucrose PBS solution overnight. Tissues were cryoembedded in mounting media (OCT), and sectioned at 20 μm thickness before being processed for immunohistochemistry with an antibody against PGP9.5 and L1CAM (Table 1). Fluorescent images were collected by confocal microscopy. Approximately 10 images per stack were flattened using the ImageJ software. Six different skin sections were measured for each footpad. The number of intraepidermal nerve fibers and nociceptive Schwann cells (PGP9.5+/DAPI+) or (L1CAM+/DAPI+) were presented as the mean number of fibers and cells per linear millimeter of epidermis. All analyses were performed by double-blinded quantification.

### Quantifications of immunolabeled LDLR and CD36

Sciatic nerve sections (14 µm) were stained with an antibody against LDLR and CD36 (Table 1). An average of 10 images were measured for every mouse, and intensity expression was calculated using ImageJ software and normalized to the DAPI intensity of each image. All analyses were performed by double-blinded quantification.

### Electron Microscopy

Mice were perfused with 20 mL 1XPBS followed by 15mL of 4% PFA (Electron Microscopy Sciences # 15172) diluted in PHEM buffer (0.1 M; pH 7.4) (Electron Microscopy Sciences # 11163). Sciatic and sural nerves were dissected and fixed in 4% PFA + 2.5% glutaraldehyde (Acros Organics, Thermo Fisher Scientific, Waltham, MA, USA) diluted in PHEM buffer overnight at 4°C. Tissues were stored in 0.5% glutaraldehyde / PHEM buffer at 4°C until use. After being washed in 0.2 M PBS buffer, the nerves were incubated with 0.5% osmic acid + 0.8% potassium hexacyanoferrate trihydrate in 0.1 M phosphate buffer for 120[minutes at room temperature. After two rinses in PHEM buffer, the nerves were dehydrated in a graded series of ethanol solutions (30-100%). The nerves were embedded in EmBed 812 using an Automated Microwave Tissue Processor for Electronic Microscopy, Leica EM AMW. Thin sections (70 nm; Leica-Reichert Ultracut E) were collected at different levels of each block. These sections were counterstained with 1.5% uranyl acetate in 70% Ethanol and lead citrate before observation using a Tecnai F20 transmission electron microscope at 120KV in the Institut des Neurosciences de Montpellier: Electronic Microscopy facilities, INSERM U 1298, Université de Montpellier, Montpellier France.

The axonal diameter was manually and blindly measured using the MyelTracer software ^50^. Mitochondria were manually and double-blindly quantified. TEM images of sciatic nerve sections were acquired at (x8,500, x10,000) magnifications of 20 randomly selected fields.

### Toluidine Blue Staining and myelinated fiber analysis

Slides were stained with 1% toluidine blue solution containing 2% borax, washed with water, and then mounted in Entellan mounting medium (EMS, Hatfield, USA). Reconstituted tiles of the whole nerves were imaged at ×63 magnification. To determine g-ratio, images were cropped into random fields. Axon and fiber (axon+myelin) diameters for at least 1162 axons per genotype (on average ≈300 axons per mouse) were calculated using MyelTracer software^50^. All quantifications were performed by an individual blinded to the animal genotype.

### Lipid Droplet Staining

For lipid droplets staining, sections were fixed in 4% paraformaldehyde, washed with 1X PBS and incubated for 45 min at 20°C with 1 µg/mL of the fluorescent neutral lipid stain Bodipy 493/503 (Life Technologies, California, USA). After washing, sections were counterstained with DAPI and mounted in vectashield (Vector Laboratories).

### Proteomics

Nerves from PCSK9 KO mice (n=4 WT mice and n=3 PCSK9 KO) were pooled until reaching 10 mg and lysed with RIPA buffer (50 mM Tris-HCl pH 7.2, 5 mM EDTA, 0.1%SDS) and Halt™ Protease Inhibitor Cocktail (Thermofisher, Massachusetts, USA) for 60 minutes at 4°C with constant shaking. Next, samples were centrifuged at 13,000 g for 5 minutes, the supernatant containing the protein extract was collected and the total protein concentration was determined (BCA Protein Assay Kit, Thermofisher). Samples were reduced with DTT (15 mM) and protein alkylation was then performed at RT, by incubating with iodoacetamide (final concentration 15 mM), for 30 minutes in the dark. Trypsin digestion (enzyme-to-protein ratio of 1:20), was performed overnight at 37°C, and the tryptic peptide digests were resuspended in 7 µL of 4% acetonitrile (ACN) in 0.1% TFA and analyzed by nano-LC using a Thermo Fisher Ultimate 3000 series NCS-3500 RS coupled with NSI-Q-Orbitrap mass spectrometer (Q Exactive Plus, Thermo Fisher Scientific, Bremen, Germany). 5 µL of sample (each sample was injected in triplicate) were separated on a LC-EASY-spray C18 column (2.6 µm, 100 Å, 75 µm[×[25 cm, Thermo Fisher Scientific). The mass spectrometry analysis was performed with the following conditions: spray voltage: 2 kV, heated capillary temperature: 275 °C, and S-lens RF level: 30%. Mass spectra were acquired with XCalibur 4.2.47 software (Thermo Fisher Scientific) and registered in data-dependent acquisition with the mass spectrometer operating in positive mode. Survey full scan mass spectra were acquired in the 350 to 2,000 m*/z* range at a resolving power of 70,000 (at m/z 400) with automatic gain control (AGC) target of 1e^6^ and maximum injection time (max IT) of 120 ms. Injection of blank (4% acetonitrile in 0.1% TFA) was performed before and after samples to prevent carry-over. The Orbitrap performance was evaluated weekly and external calibration of the mass spectrometer was performed prior to analysis with an LTQ ESI positive ion calibration solution (Pierce).

### Protein identification and quantification

Raw mass spectrometry data were automatically processed using Proteome discoverer software (version 2.2.2.2.0, Thermo Fisher Scientific) for protein identification and quantification. MS and MS/MS spectra were searched against the UniProt mouse reference proteome database with canonical and isoform sequences. The database search was performed with the following parameters: oxidized methionine and protein N-terminal acetylation were set as variable modifications, and Cysteine carbamidomethylation was set as a fixed modification. Trypsin was set as enzyme-specific and two missed cleavages were allowed. A mass tolerance of 10 ppm was used for precursor ions and 0.02 Da for-product ions. The false-discovery rate (FDR) was fixed at 1% at the level of proteins and peptides using a target-reversed decoy database search strategy. A minimum of one unique peptide sequence with a Sequest score (Xcorr)[≥[2 was used. For peptides with an Xcorr[<[2, identification was confirmed by manual interpretation of the corresponding MS/MS spectrum.

### Statistical analysis

Graph Pad-Prism 8 (GraphPad Software Inc., San Diego, CA, USA) was used for graphic presentation and statistical analysis. Results are expressed as the mean per group ± SD. We considered *p<0.05, **p<0.01, ***p<0.001 and **** *P* <0.0001 as significant. For a two-group comparison, a T-test was used. For multiple comparisons, a one-way ANOVA followed by Tukey’s multiple comparisons test was used. For the integration of the proteome for functional analysis, protein abundance was calculated with the generation of spectral feature by the node Feature Finder Multiplex followed by PIA-assisted FDR-multiple scores estimation and filtering (combined FDR score<0.01), their ID mapping and combination with peptide IDs, their subsequent alignment, grouping, and normalization. DAVID (The Database for Annotation, Visualization, and Integrated Discovery) platform was used for gene ontology (GO) enrichment analysis. GO was performed in our protein LFQ dataset and results for significant terms associated with the gene set were selected based on the fold discovery rate FDR<0.05.

## Study approval

All *in vivo* experiments were conducted in accordance with the European Community Guidelines for the Use of Animals in Research (86/609/EEC and 2010/63/EU) and approved by the local Ethics Committee for animal experimentation (APAFIS#3209-2015111215451823v2).

## Data availability

The corresponding author will provide all requested materials, data sets, and protocols, without restriction, upon request.

## Author contributions

AKJ and SB designed the studies. AKJ, APR, GR, BV, CP, MB and PR carried out the experiments. AKJ, GL and SB interpreted the data. AKJ, OM, GL and SB drafted the manuscript, and participated in its writing and editing.

## Acknowledgments

AKJ is the recipient of a PhD scholarship from La Fondation pour la Recherche Médicale (Paris, France) and from the European Union and Region Réunion (Saint-Denis, Réunion); GL is the recipient of a project grant KRINGLE2 ANR-20-CE14-0009 funded by the Agence Nationale de la Recherche (Paris, France); SB is the recipient of a program grant ATIP-AVENIR funded by Inserm (Paris, France). We would like to acknowledge Alexandre Pattyn (Institut des Neurosciences de Montpellier) for discussions and helpful comments on this study, Chantal Cazevieille (Institut des Neurosciences de Montpellier) for her help in electron microscopy analysis.

## Abbreviations

ApoER2: Apolipoprotein E receptor 2
ASCVD: Atherosclerotic cardiovascular diseases
CD36: Cluster of differentiation 36
DRG: Dorsal root ganglia
IENFD: Intraepidermal nerve fiber density
LDLR: Low-density lipoprotein receptor
LRP1: LDLR-related protein 1
NARC-1: Neural Apoptosis-Regulated Convertase-1
NCV: Nerve conduction velocity
PCSK9: Proprotein convertase subtilisin/kexin type 9
PNS: Peripheral nervous system
SCs: Schwann cells
VLDLR: Very-low-density lipoprotein receptor

---

**Supplemental figure 1.**
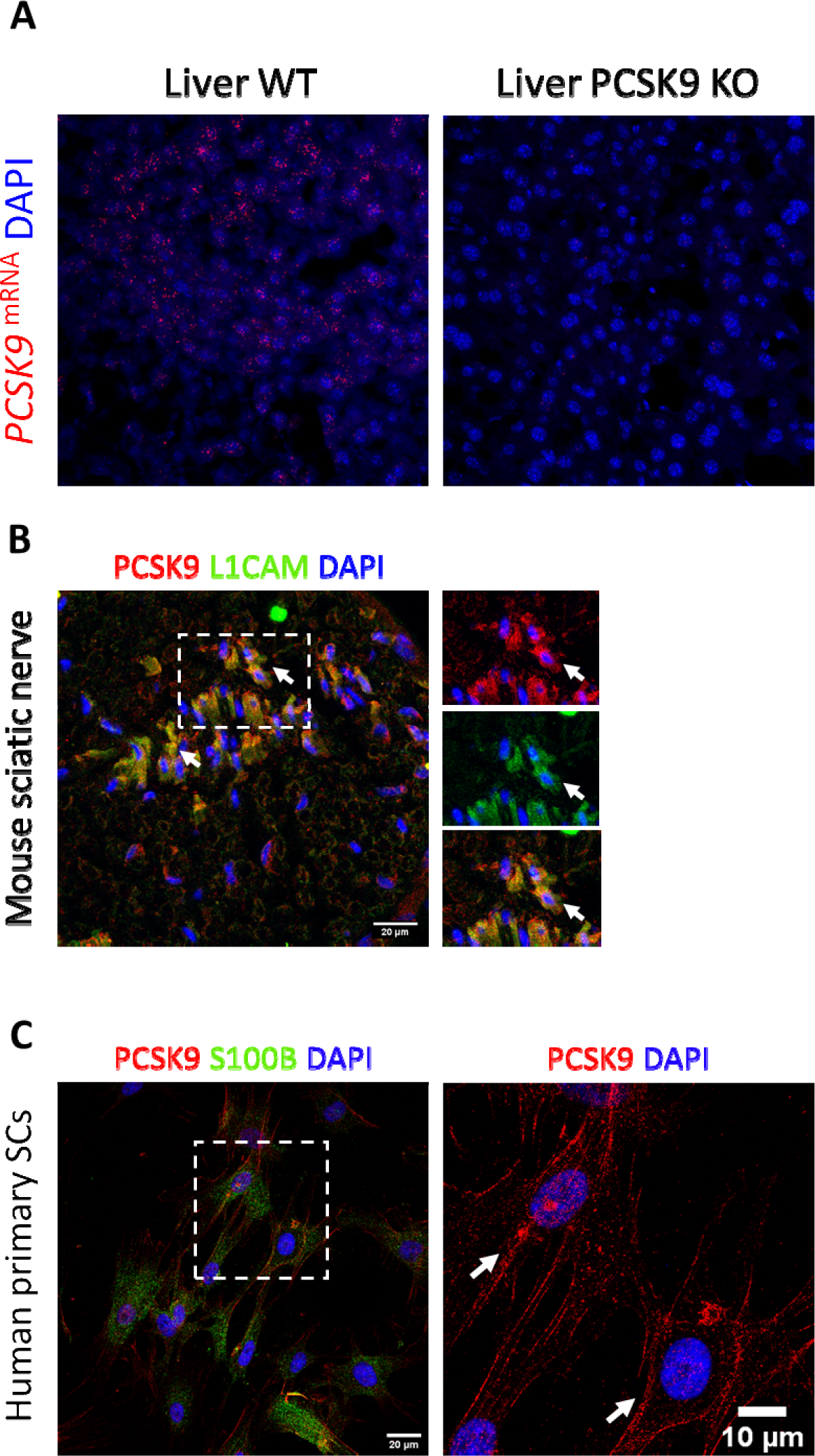
**(A)** PCSK9 mRNA expression in mouse liver of control (WT) and PCSK9 KO mice. **(B)** PCSK9 expression in Schwann cell subtypes in mouse sciatic nerve, showing colocalization of PCSK9 with L1CAM positive non–myelinating Schwann cells. **(C)** Expression of PCSK9 in Human primary Schwann cells culture (arrows: PCSK9 expression).

**Supplemental figure 2.**
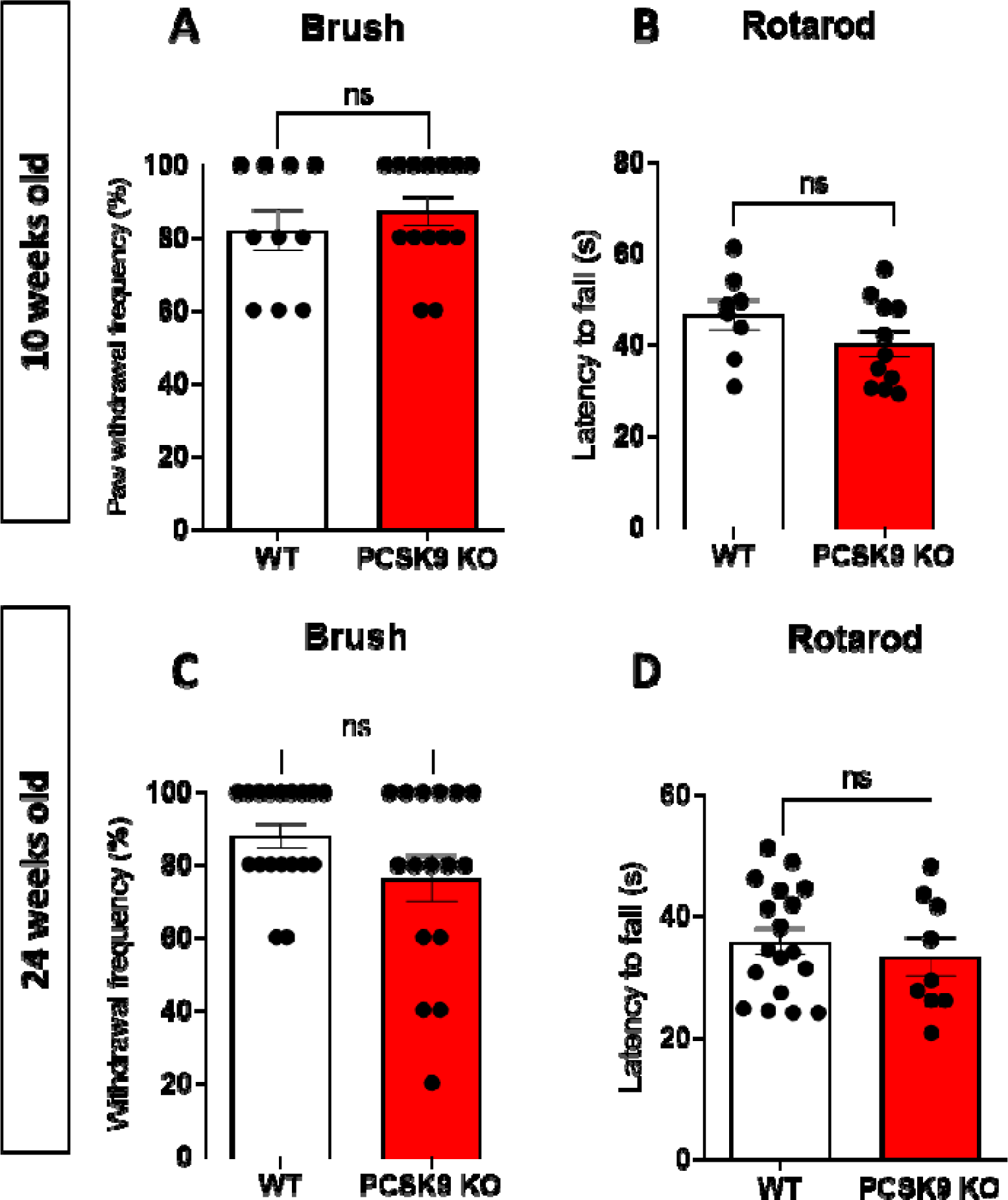
**(A)** Dynamic touch sensation in 10-week-old WT (n=10) and PCSK9 KO (n=14) mice with the brush test. **(B)** Gross motor coordination in 10-week-old WT (n=8) and PCSK9 KO (n=11) mice with the rotarod test. **(C)** Dynamic touch sensation in 24-week-old WT (n=18) and PCSK9 KO (n=16) mice with the brush test. **(D)** Gross motor coordination in 24-week-old WT (n=18) and PCSK9 KO (n=9) mice with the rotarod test. Data are represented as mean ± SEM and statistically analyzed by unpaired t test. *P* > 0.05 (ns).

**Supplemental figure 3.**
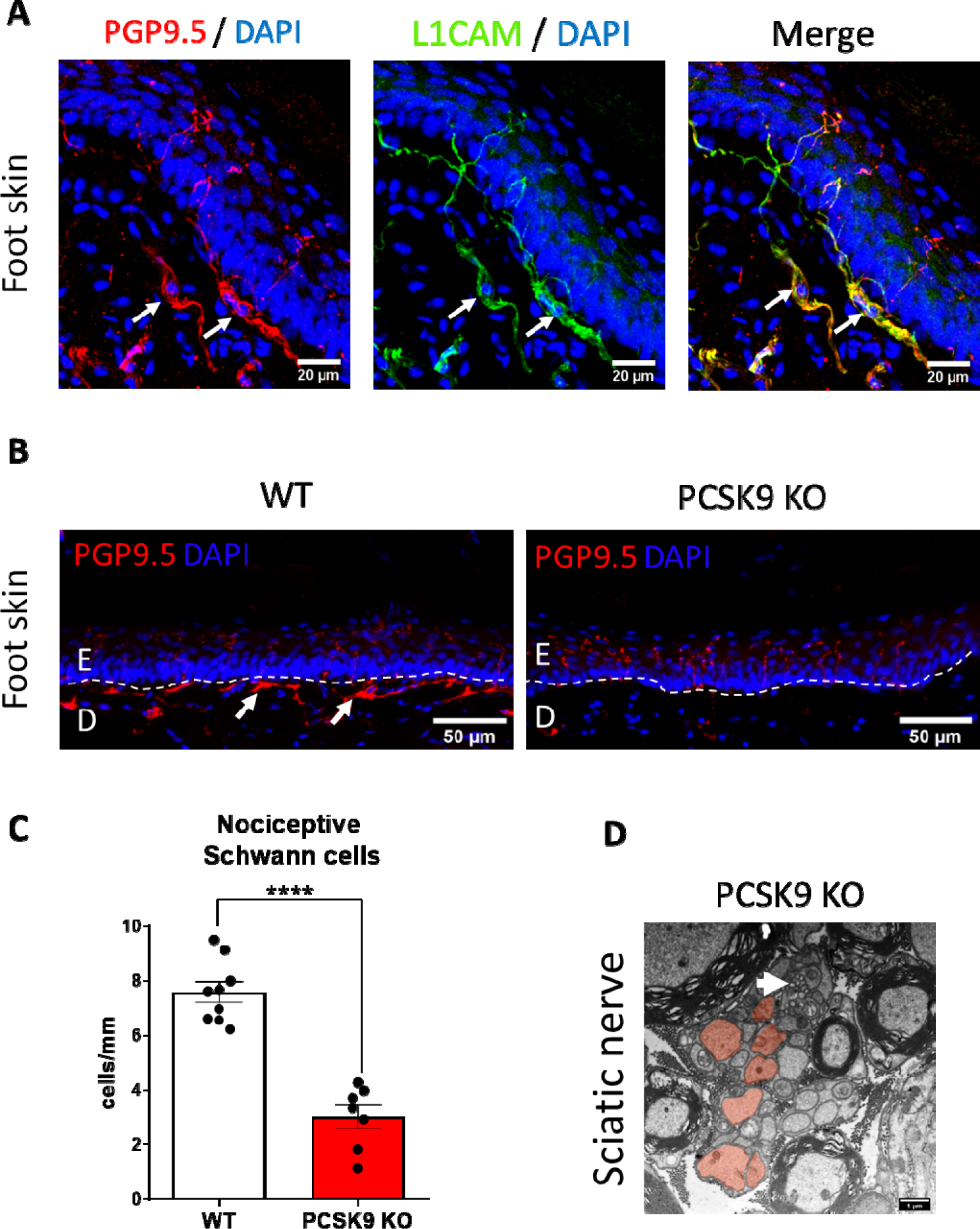
**(A)** Images of terminal nociceptive Schwann cells with colocalization between PGP9.5 and L1CAM. **(B)** Representative images of nociceptive Schwann cells identified by immunostaining against PGP9.5/DAPI in the footpad of WT and PCSK9 KO mice. **(C)** Quantification of the number of nociceptive Schwann cells in WT (n=9) and PCSK9 KO (n=7) mice. **(D)** Electron microscopy picture of sciatic nerve showing structural abnormalities in axonal circularity in PCSK9 KO mice (red highlight). Arrow shows vacuoles inside the nerve. Data are represented as mean ± SEM and statistically analyzed by unpaired t test. \*\*\*\**P*≤0.0001.

**Supplemental figure 4.**
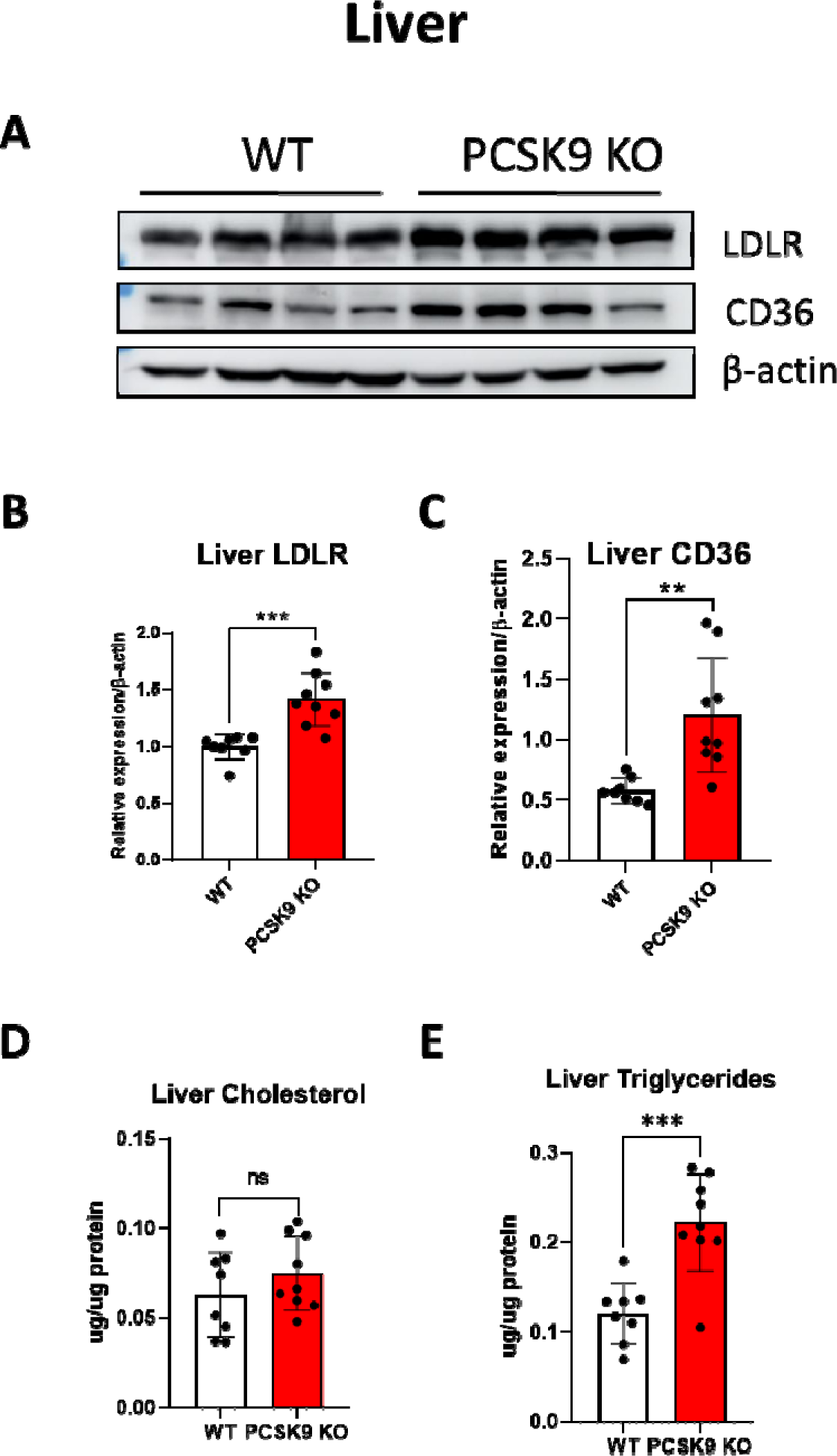
**(A)** LDLR and CD36 expression in the liver by immunoblotting. **(B)** Quantification of LDLR liver expression in 24-week-old WT (n=9) and PCSK9 KO (n=9). **(C)** Quantification of CD36 liver expression in 24-week-old WT (n=8) and PCSK9 KO (n=9). **(D)** Cholesterol levels in the liver of 24-week-old WT (n=8) and PCSK9 KO (n=9) mice. **(E)** Triglyceride levels in the liver of 24-week-old WT (n=8) et PCSK9 KO (n=9) mice. Data are represented as mean ± SEM and statistically analyzed by unpaired t test. *P* > 0.05 (ns), \*\**P* < 0.01, and \*\*\**P* < 0.001.

**Supplemental figure 5.**
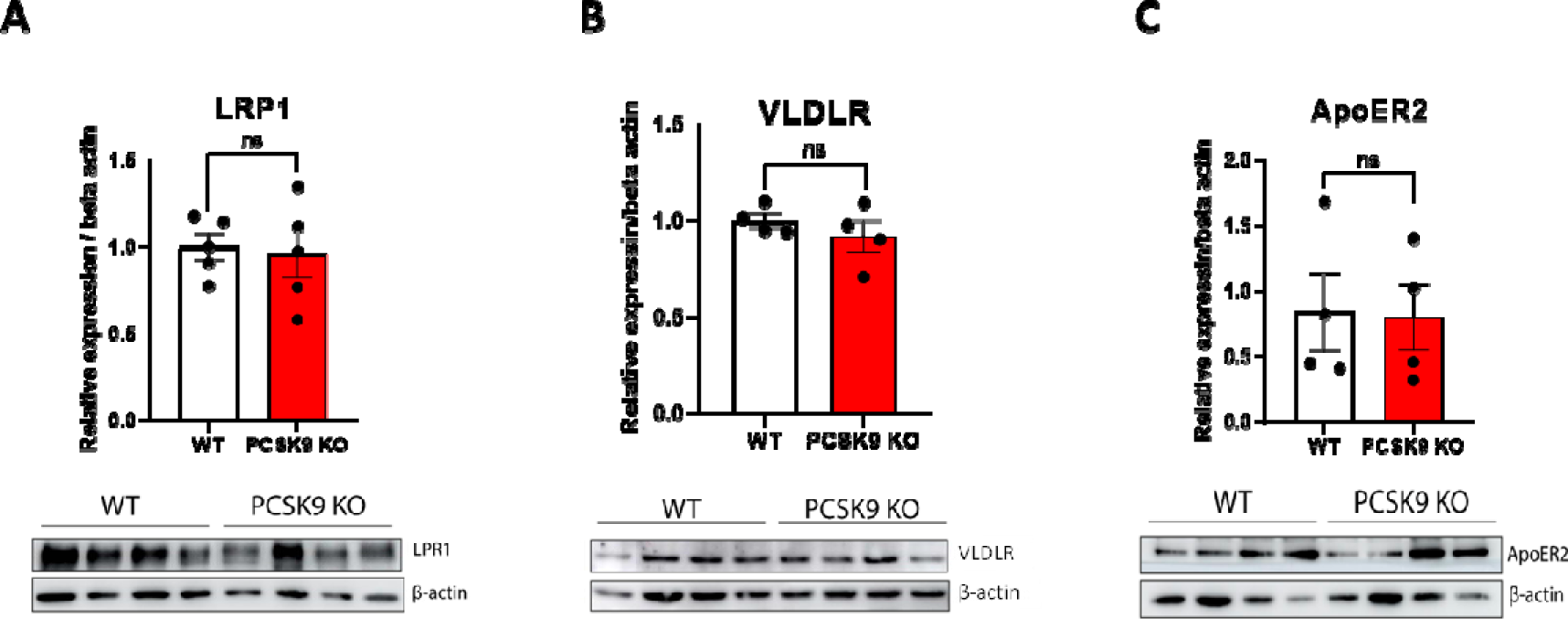
**(A)** Quantification of LRP1 expression in the sciatic nerve of 24-week-old WT (n=5) and PCSK9 KO (n=5) mice. **(B)** Quantification of VLDLR expression in the sciatic nerve of 24-week-old WT (n=5) and PCSK9 KO (n=5). **(C)** Quantification of ApoER2 expression in the sciatic nerve of 24-week-old WT (n=5) and PCSK9 KO (n=5). Data are represented as mean ± SEM and statistically analyzed by unpaired t test. *P* > 0.05 (ns).

**Supplemental figure 6.**
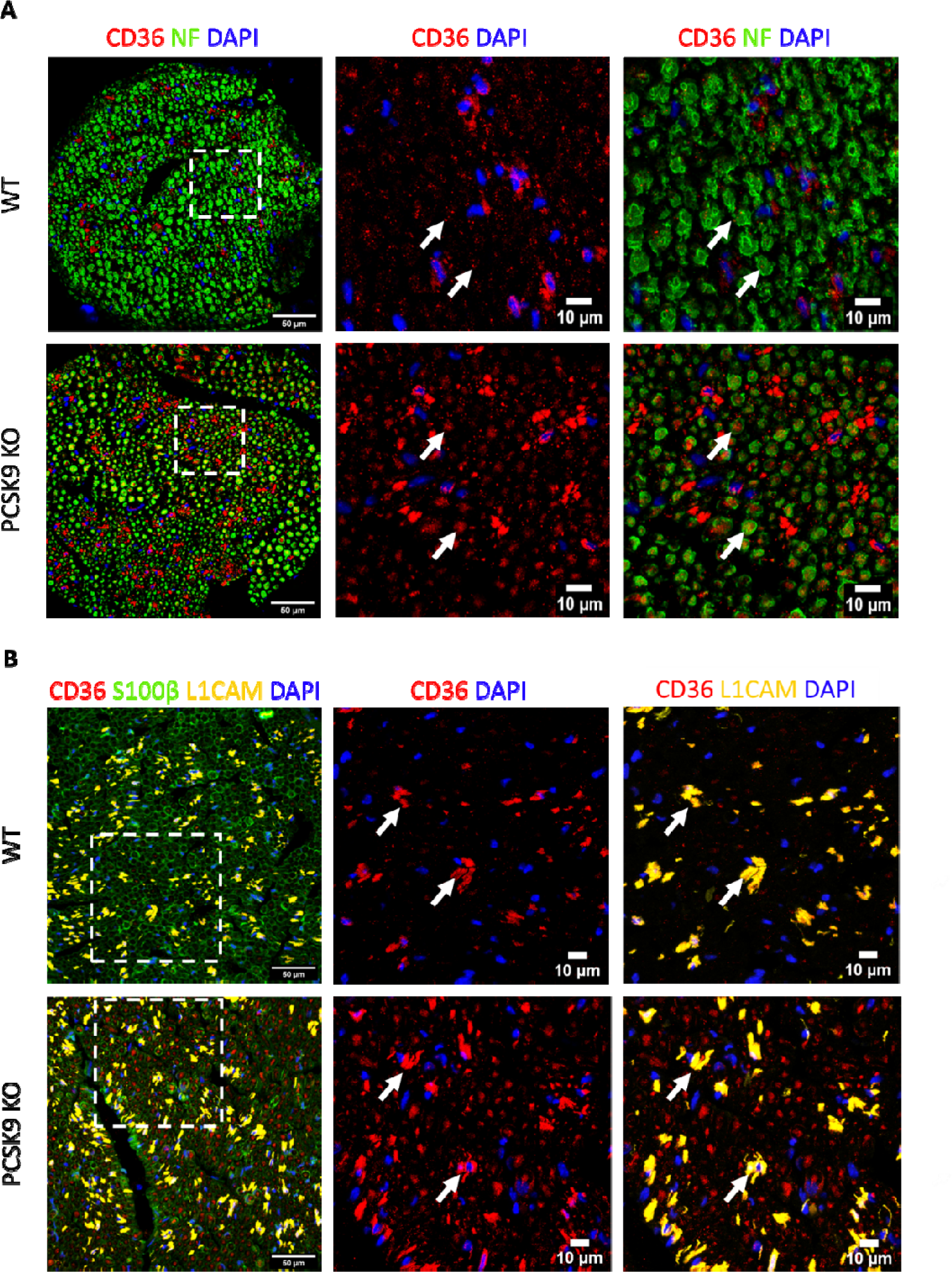
**(A)** Transverse section of mouse sciatic nerve of 24-week-old WT and PCSK9 KO labeled with an antibody against CD36 (red) and neurofilament (NF) (green) as well as nuclear DAPI staining (blue). **(B)** Triple immunostaining of transverse section mouse sciatic nerve of 24-week-old WT and PCSK9 KO with antibody against CD36 (red), S100B a pan-Schwann cell marker (green) and L1CAM a non-myelinating Schwann cell marker (yellow) as well as nuclear DAPI staining (nuclear staining blue). (Arrows show the colocalization of CD36 with the different markers).

**Supplemental figure 7.**
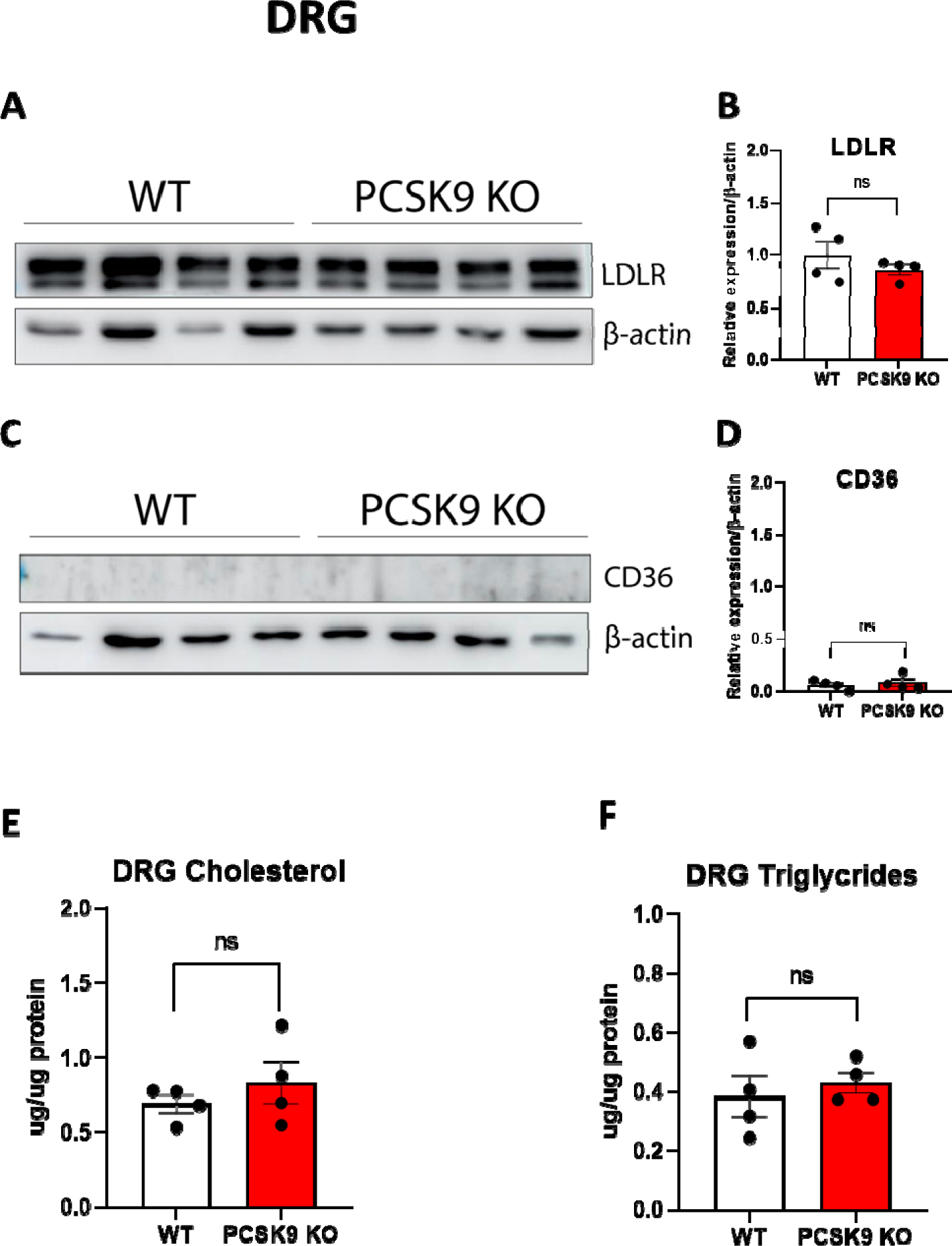
**(A)** Immunoblot of LDLR expression in DRG of 24-week-old WT and PCSK9 KO mice. **(B)** Quantification of LDLR expression level in DRG of WT (n=4) and PCSK9 KO (n=4) mice. **(C)** Immunoblot of CD36 expression in DRG of 24-week-old WT and PCSK9 KO mice. **(D)** Quantification of CD36 expression level in DRG of WT (n=4) and PCSK9 KO (n=4) mice. **(E)** Quantification of cholesterol level in DRG of 24 weeks WT (n=4) and PCSK9 KO (n=4) mice. **(F)** Quantification of triglyceride levels in DRG of 24-weeks-old WT (n=4) and PCSK9 KO (n=4) mice. Data are represented as mean ± SEM and statistically analyzed by unpaired t test. *P* > 0.05 (ns).

**Supplemental figure 8.**
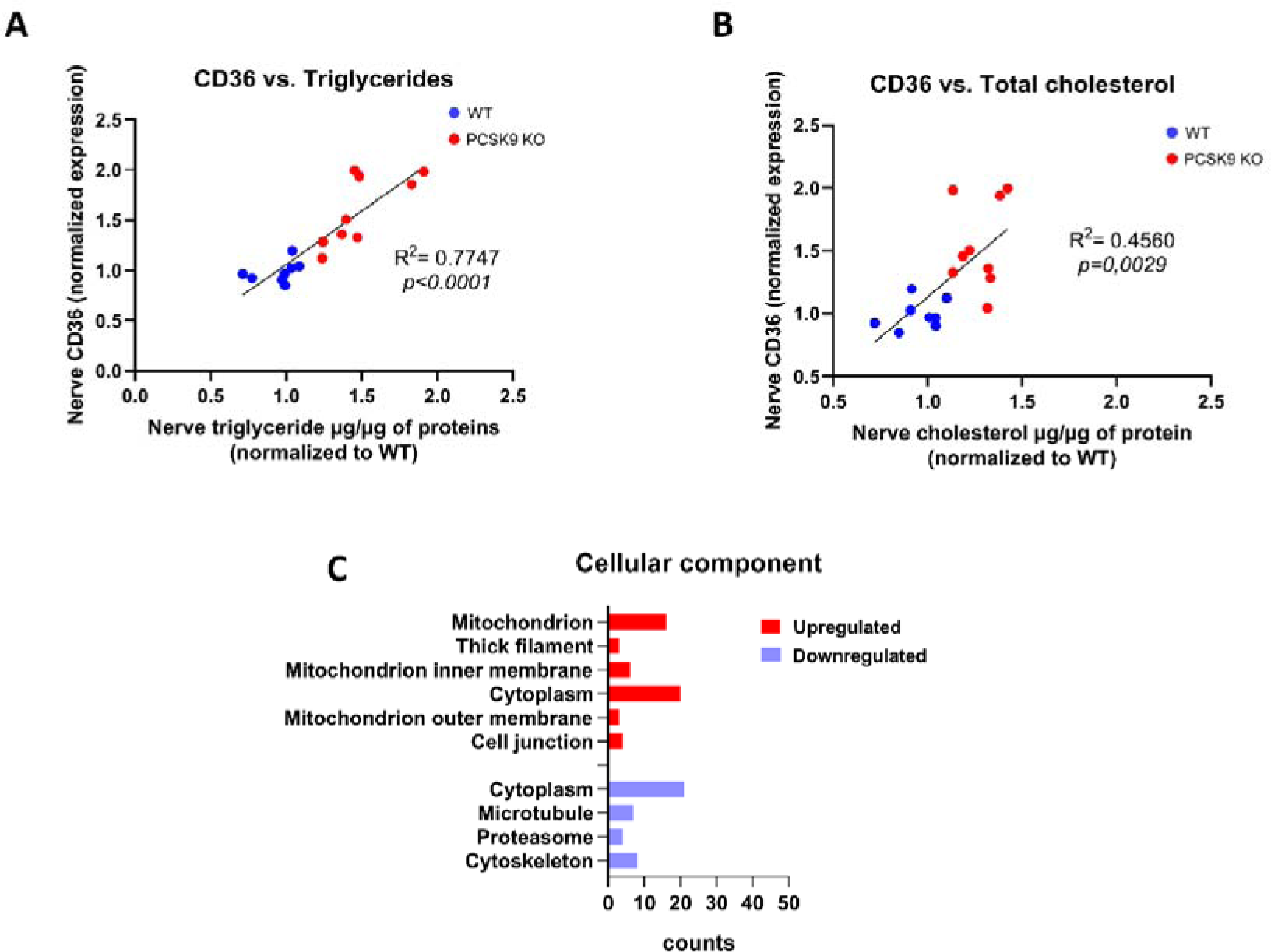
**(A)** Scatter plot displaying the correlation between nerve CD36 expression and nerve triglyceride levels in WT (n=8) and PCSK9 KO (n=9). **(B)** Scatter plot displaying the correlation between nerve CD36 expression and nerve cholesterol levels in WT (n=8) and PCSK9 KO (n=9). **(C)** Gene ontology annotation of the top upregulated (red) and downregulated (blue) in terms of molecular function and (D) cellular component sorted by P-value.

## References

1. Seidah NG, Garçon D. Expanding Biology of PCSK9: Roles in Atherosclerosis and Beyond. Curr Atheroscler Rep. 2022;24(10):821–830. doi:10.1007/s11883-022-01057-z

2. Cohen, Boerwinkle E, Mosley TH, Hobbs HH. Sequence variations in PCSK9, low LDL, and protection against coronary heart disease. N Engl J Med. 2006;354(12):1264–1272. doi:10.1056/NEJMoa054013

3. Abifadel M, Varret M, Rabès JP, et al. Mutations in PCSK9 cause autosomal dominant hypercholesterolemia. Nature Genetics. 2003;34(2):154–156. doi:10.1038/ng1161

4. Sabatine MS, Giugliano RP, Keech AC, et al. Evolocumab and Clinical Outcomes in Patients with Cardiovascular Disease. New England Journal of Medicine. 2017;376(18):1713–1722. doi:10.1056/NEJMoa1615664

5. Schwartz GG, Steg PG, Szarek M, et al. Alirocumab and Cardiovascular Outcomes after Acute Coronary Syndrome. New England Journal of Medicine. 2018;379(22):2097–2107. doi:10.1056/NEJMoa1801174

6. Zaid A, Roubtsova A, Essalmani R, et al. Proprotein convertase subtilisin/kexin type 9 (PCSK9): Hepatocyte-specific low-density lipoprotein receptor degradation and critical role in mouse liver regeneration. Hepatology. 2008;48(2):646–654. doi:10.1002/hep.22354

7. Seidah NG, Benjannet S, Wickham L, et al. The secretory proprotein convertase neural apoptosis-regulated convertase 1 (NARC-1): liver regeneration and neuronal differentiation. Proc Natl Acad Sci U S A. 2003;100(3):928–933. doi:10.1073/pnas.0335507100

8. Jaafar AK, Techer R, Chemello K, Lambert G, Bourane S. PCSK9 and the nervous system: a no-brainer? Journal of Lipid Research. 2023;64(9). doi:10.1016/j.jlr.2023.100426

9. O’Connell EM, Lohoff FW. Proprotein Convertase Subtilisin/Kexin Type 9 (PCSK9) in the Brain and Relevance for Neuropsychiatric Disorders. Front Neurosci. 2020;14:609. doi:10.3389/fnins.2020.00609

10. Da Dalt L, Ruscica M, Bonacina F, et al. PCSK9 deficiency reduces insulin secretion and promotes glucose intolerance: the role of the low-density lipoprotein receptor. Eur Heart J. 2019;40(4):357–368. doi:10.1093/eurheartj/ehy357

11. Poitelon Y, Kopec AM, Belin S. Myelin Fat Facts: An Overview of Lipids and Fatty Acid Metabolism. Cells. 2020;9(4):812. doi:10.3390/cells9040812

12. Fu Q, Goodrum JF, Hayes C, Hostettler JD, Toews AD, Morell P. Control of cholesterol biosynthesis in Schwann cells. J Neurochem. 1998;71(2):549–555. doi:10.1046/j.1471-4159.1998.71020549.x

13. Montani L, Pereira JA, Norrmén C, et al. De novo fatty acid synthesis by Schwann cells is essential for peripheral nervous system myelination. J Cell Biol. 2018;217(4):1353–1368. doi:10.1083/jcb.201706010

14. Sánchez-Alegría K, Bastián-Eugenio CE, Vaca L, Arias C. Palmitic acid induces insulin resistance by a mechanism associated with energy metabolism and calcium entry in neuronal cells. FASEB J. 2021;35(7):e21712. doi:10.1096/fj.202100243R

15. Rothe T, Müller HW. Uptake of endoneurial lipoprotein into Schwann cells and sensory neurons is mediated by low density lipoprotein receptors and stimulated after axonal injury. J Neurochem. 1991;57(6):2016–2025. doi:10.1111/j.1471-4159.1991.tb06417.x

16. Goodrum JF, Fowler KA, Hostettler JD, Oews ADT. Peripheral nerve regeneration and cholesterol reutilization are normal in the low-density lipoprotein receptor knockout mouse. Journal of Neuroscience Research. 2000;59(4):581–586. doi:10.1002/(SICI)1097-4547(20000215)59:4<581::AID-JNR14>3.0.CO;2-P

17. Orita S, Henry K, Mantuano E, et al. Schwann cell LRP1 regulates remak bundle ultrastructure and axonal interactions to prevent neuropathic pain. J Neurosci. 2013;33(13):5590–5602. doi:10.1523/JNEUROSCI.3342-12.2013

18. Pasten C, Cerda J, Jausoro I, Court FA, Cáceres A, Marzolo MP. ApoER2 and Reelin are expressed in regenerating peripheral nerve and regulate Schwann cell migration by activating the Rac1 GEF protein, Tiam1. Mol Cell Neurosci. 2015;69:1–11. doi:10.1016/j.mcn.2015.09.004

19. Toma JS, Karamboulas K, Carr MJ, et al. Peripheral Nerve Single-Cell Analysis Identifies Mesenchymal Ligands that Promote Axonal Growth. eNeuro. 2020;7(3):ENEURO.0066-20.2020. doi:10.1523/ENEURO.0066-20.2020

20. Abdo H, Calvo-Enrique L, Lopez JM, et al. Specialized cutaneous Schwann cells initiate pain sensation. Science. 2019;365(6454):695–699. doi:10.1126/science.aax6452

21. Ioghen O, Manole E, Gherghiceanu M, et al. Non-Myelinating Schwann Cells in Health and Disease. In: Demyelination Disorders. IntechOpen; 2020. doi:10.5772/intechopen.91930

22. Lebeau PF, Byun JH, Platko K, et al. Pcsk9 knockout exacerbates diet-induced non-alcoholic steatohepatitis, fibrosis and liver injury in mice. JHEP Rep. 2019;1(6):418–429. doi:10.1016/j.jhepr.2019.10.009

23. Arcourt A, Gorham L, Dhandapani R, et al. Touch Receptor-Derived Sensory Information Alleviates Acute Pain Signaling and Fine-Tunes Nociceptive Reflex Coordination. Neuron. 2017;93(1):179–193. doi:10.1016/j.neuron.2016.11.027

24. Dubin AE, Patapoutian A. Nociceptors: the sensors of the pain pathway. J Clin Invest. 2010;120(11):3760–3772. doi:10.1172/JCI42843

25. Benjannet S, Rhainds D, Essalmani R, et al. NARC-1/PCSK9 and Its Natural Mutants: ZYMOGEN CLEAVAGE AND EFFECTS ON THE LOW DENSITY LIPOPROTEIN (LDL) RECEPTOR AND LDL CHOLESTEROL*. Journal of Biological Chemistry. 2004;279(47):48865–48875. doi:10.1074/jbc.M409699200

26. Zhang DW, Lagace TA, Garuti R, et al. Binding of Proprotein Convertase Subtilisin/Kexin Type 9 to Epidermal Growth Factor-like Repeat A of Low Density Lipoprotein Receptor Decreases Receptor Recycling and Increases Degradation*. Journal of Biological Chemistry. 2007;282(25):18602–18612. doi:10.1074/jbc.M702027200

27. Schmidt RJ, Beyer TP, Bensch WR, et al. Secreted proprotein convertase subtilisin/kexin type 9 reduces both hepatic and extrahepatic low-density lipoprotein receptors in vivo. Biochem Biophys Res Commun. 2008;370(4):634–640. doi:10.1016/j.bbrc.2008.04.004

28. Liu M, Wu G, Baysarowich J, et al. PCSK9 is not involved in the degradation of LDL receptors and BACE1 in the adult mouse brain. Journal of Lipid Research. 2010;51(9):2611–2618. doi:10.1194/jlr.M006635

29. Pärn A, Olsen D, Tuvikene J, et al. PCSK9 deficiency alters brain lipid composition without affecting brain development and function. Front Mol Neurosci. 2022;15:1084633. doi:10.3389/fnmol.2022.1084633

30. Mayer G, Poirier S, Seidah NG. Annexin A2 is a C-terminal PCSK9-binding protein that regulates endogenous low density lipoprotein receptor levels. J Biol Chem. 2008;283(46):31791–31801. doi:10.1074/jbc.M805971200

31. Seidah NG, Poirier S, Denis M, et al. Annexin A2 Is a Natural Extrahepatic Inhibitor of the PCSK9-Induced LDL Receptor Degradation. PLoS One. 2012;7(7):e41865. doi:10.1371/journal.pone.0041865

32. Ly K, Essalmani R, Desjardins R, Seidah NG, Day R. An Unbiased Mass Spectrometry Approach Identifies Glypican-3 as an Interactor of Proprotein Convertase Subtilisin/Kexin Type 9 (PCSK9) and Low Density Lipoprotein Receptor (LDLR) in Hepatocellular Carcinoma Cells. J Biol Chem. 2016;291(47):24676–24687. doi:10.1074/jbc.M116.746883

33. Jang HD, Lee SE, Yang J, et al. Cyclase-associated protein 1 is a binding partner of proprotein convertase subtilisin/kexin type-9 and is required for the degradation of low-density lipoprotein receptors by proprotein convertase subtilisin/kexin type-9. Eur Heart J. 2020;41(2):239–252. doi:10.1093/eurheartj/ehz566

34. Demers A, Samami S, Lauzier B, et al. PCSK9 Induces CD36 Degradation and Affects Long-Chain Fatty Acid Uptake and Triglyceride Metabolism in Adipocytes and in Mouse Liver. Arteriosclerosis, Thrombosis, and Vascular Biology. 2015;35(12):2517–2525. doi:10.1161/ATVBAHA.115.306032

35. Da Dalt L, Castiglioni L, Baragetti A, et al. PCSK9 deficiency rewires heart metabolism and drives heart failure with preserved ejection fraction. Eur Heart J. 2021;42(32):3078–3090. doi:10.1093/eurheartj/ehab431

36. Byun JH, Lebeau PF, Platko K, et al. Inhibitory Antibodies against PCSK9 Reduce Surface CD36 and Mitigate Diet-Induced Renal Lipotoxicity. Kidney360. 2022;3(8):1394–1410. doi:10.34067/KID.0007022021

37. Cartelli D, Cavaletti G, Lauria G, Meregalli C. Ubiquitin Proteasome System and Microtubules Are Master Regulators of Central and Peripheral Nervous System Axon Degeneration. Cells. 2022;11(8):1358. doi:10.3390/cells11081358

38. VerPlank JJS, Lokireddy S, Feltri ML, Goldberg AL, Wrabetz L. Impairment of protein degradation and proteasome function in hereditary neuropathies. Glia. 2018;66(2):379–395. doi:10.1002/glia.23251

39. Edwards JL, Quattrini A, Lentz SI, et al. Diabetes regulates mitochondrial biogenesis and fission in mouse neurons. Diabetologia. 2010;53(1):160–169. doi:10.1007/s00125-009-1553-y

40. Vincent AM, Edwards JL, McLean LL, et al. Mitochondrial biogenesis and fission in axons in cell culture and animal models of diabetic neuropathy. Acta Neuropathol. 2010;120(4):477–489. doi:10.1007/s00401-010-0697-7

41. Franco DC, Neupane N, Riaz M, Mohammadzadeh S, Sachmechi I. Chronic Inflammatory Demyelinating Polyradiculoneuropathy Association With Low Cholesterol Levels: A Case Report in a Patient Taking PCSK9 Inhibitor. Journal of Neurology Research. 2019;9(4-5):72–74. doi:10.14740/jnr.v9i4-5.552

42. Robinson JG, Farnier M, Krempf M, et al. Efficacy and safety of alirocumab in reducing lipids and cardiovascular events. N Engl J Med. 2015;372(16):1489–1499. doi:10.1056/NEJMoa1501031

43. Ray KK, Wright RS, Kallend D, et al. Two Phase 3 Trials of Inclisiran in Patients with Elevated LDL Cholesterol. N Engl J Med. 2020;382(16):1507–1519. doi:10.1056/NEJMoa1912387

44. Eto M, Yoshikawa H, Fujimura H, et al. The role of CD36 in peripheral nerve remyelination after crush injury. The European journal of neuroscience. 2003;17:2659–2666. doi:10.1046/j.1460-9568.2003.02711.x

45. Eto M, Sumi H, Fujimura H, Yoshikawa H, Sakoda S. Pioglitazone promotes peripheral nerve remyelination after crush injury through CD36 upregulation. J Peripher Nerv Syst. 2008;13(3):242–248. doi:10.1111/j.1529-8027.2008.00183.x

46. Roubtsova A, Chamberland A, Marcinkiewicz J, et al. PCSK9 deficiency unmasks a sex- and tissue-specific subcellular distribution of the LDL and VLDL receptors in mice. J Lipid Res. 2015;56(11):2133–2142. doi:10.1194/jlr.M061952

47. Roubtsova A, Garçon D, Lacoste S, et al. PCSK9 deficiency results in a specific shedding of excess LDLR in female mice only: Role of hepatic cholesterol. Biochim Biophys Acta Mol Cell Biol Lipids. 2022;1867(12):159217. doi:10.1016/j.bbalip.2022.159217

48. Tao Y. Isolation and culture of Schwann cells. Methods Mol Biol. 2013;1018:93–104. doi:10.1007/978-1-62703-444-9_9

49. Bonin RP, Bories C, De Koninck Y. A Simplified Up-Down Method (SUDO) for Measuring Mechanical Nociception in Rodents Using von Frey Filaments. Mol Pain. 2014;10:1744-8069-10-26. doi:10.1186/1744-8069-10-26

50. Kaiser T, Allen HM, Kwon O, et al. MyelTracer: A Semi-Automated Software for Myelin g-Ratio Quantification. eNeuro. 2021;8(4):ENEURO.0558-20.2021. doi:10.1523/ENEURO.0558-20.2021

